# Enumerating the chemical exposome using in-silico transformation analysis – an example using insecticides

**DOI:** 10.1101/2025.09.04.674275

**Authors:** Mannivannan Jothiramajayam, Dinesh Barupal

**Affiliations:** Integrated Data Science Laboratory for Metabolomics and Exposomics, Department of Environmental Medicine, Icahn School of Medicine at Mount Sinai, New York, USA

**Keywords:** Metabolomics, Exposome, Computational Chemistry, In-silico metabolism, ISTA, RDChiral, RXNMapper, Rxn-INSIGHT, reaction databases, Insecticides

## Abstract

The exposome encompasses a vast chemical space that can originate from the consumer industry and environmental sources. Once these chemicals enter into cells (human or other organisms), they can be also transformed into products that differ in terms of toxicity and health effects. Recent developments in machine learning methods and chemical data science resources have enabled the in-silico enumeration of transformation products. Here, we report an integrated workflow of these existing resources (RXNMmapper, Rxn-INSIGHT and RDChiral) to enumerate the transformation product for a chemical. We have generated a large library of reaction templates from > 80,000 reactions sourced from the PubChem database. Utility of the reaction screening and transformation enumeration workflow has been demonstrated for insecticide structures (n=181), yielding 19,392 unique transformation products. Use of filters and ranking by thermodynamic stability, species association, enzyme information and ADMET properties, can prioritize the products relevant for different contexts. Many of these products have PubChem entries but have not yet been linked with the parent compounds. The presented approach can be helpful in enumerating relevant chemical space for exposome using known reaction chemistry, which may ultimately contribute to expanding of the exposomic knowledgebases.

## Introduction

The chemical exposome represents the totality of an individual’s chemical exposures from all sources throughout their lifespan^1, 2^. This concept provides a comprehensive framework for understanding how the environment influences human health and diseases^3-7^. Human biomonitoring programs using targeted and untargeted assays have been instrumental in quantifying and cataloguing the body burden of chemical exposome^8-10^. These efforts have demonstrated widespread exposure to various classes of chemicals, including pesticides, food-additives, industrial contaminants, and environmental toxins. Once these chemicals enter the human body, they may get transformed by xenobiotic metabolism and other enzymatic reactions, generating additional toxic chemicals^11-14^. Targeted and untargeted biomonitoring assays rely on existing chemical exposome databases such as the Blood Exposome Database^15^ for finding high priority chemicals that are expected in a human biospecimen. Although, the database catalogues over 50,000 chemicals, the vast space of the transformation products, which is the “dark matter” of the chemical exposome, remains largely unexplored.

To expand the chemical exposome, we can use computational methods^16^ to first enumerate the transformation products for an exposure chemical. Such in-silico approach is often used in drug discovery pipelines to identify potential drug metabolites for preclinical toxicity studies^17, 18^. Similarly, in-silico transformation analysis (ISTA) for exposome chemicals can contribute to discoveries of new xenobiotic metabolic pathways and generate new hypotheses regarding how chemical exposome is processed inside the body, as well as how chemicals are transformed by non-human reaction chemistries.

A range of existing computational resources can be integrated into a framework for ISTA, including PubChem^19^ for chemical information including reaction data, RXNMapper^20^ for computing atomic mapping of reaction and RDChiral^21^ for extracting reaction templates. A reaction template captures the transformation patterns within an atom-mapped reaction which can be applied to a chemical structure to enumerate a new product. We can enumerate hundreds of new in-silico chemicals by utilizing the large-volume of reaction data from PubChem. Many of these chemicals might be already known structures but not yet linked with a known reaction.

In this study, we have utilized the PubChem reaction data to create a large-database of reaction templates and then we have established an in-silico transformation analysis workflow (Figure 1) using existing methods and applied it to known insecticide structures. Insecticides, though developed for agricultural use, have led to human exposure via food, environment and occupation. This requires biomonitoring of both parent compounds and their biotransformation products for comprehensive risk assessment and to associate exposure with potential adverse health outcomes. By integrating in-silico enumeration with databases and literature-backed validation and computational assessment of thermodynamic stability and ADME-Tox (Absorption, Distribution, Metabolism, Excretion, and Toxicity) properties, this study presents a useful framework for enumerating and ranking products for a chemical group relevant to the human exposome. These enumerated products cover already known products but also new hypotheses for future experimental and computational analysis.

**Figure 1.**
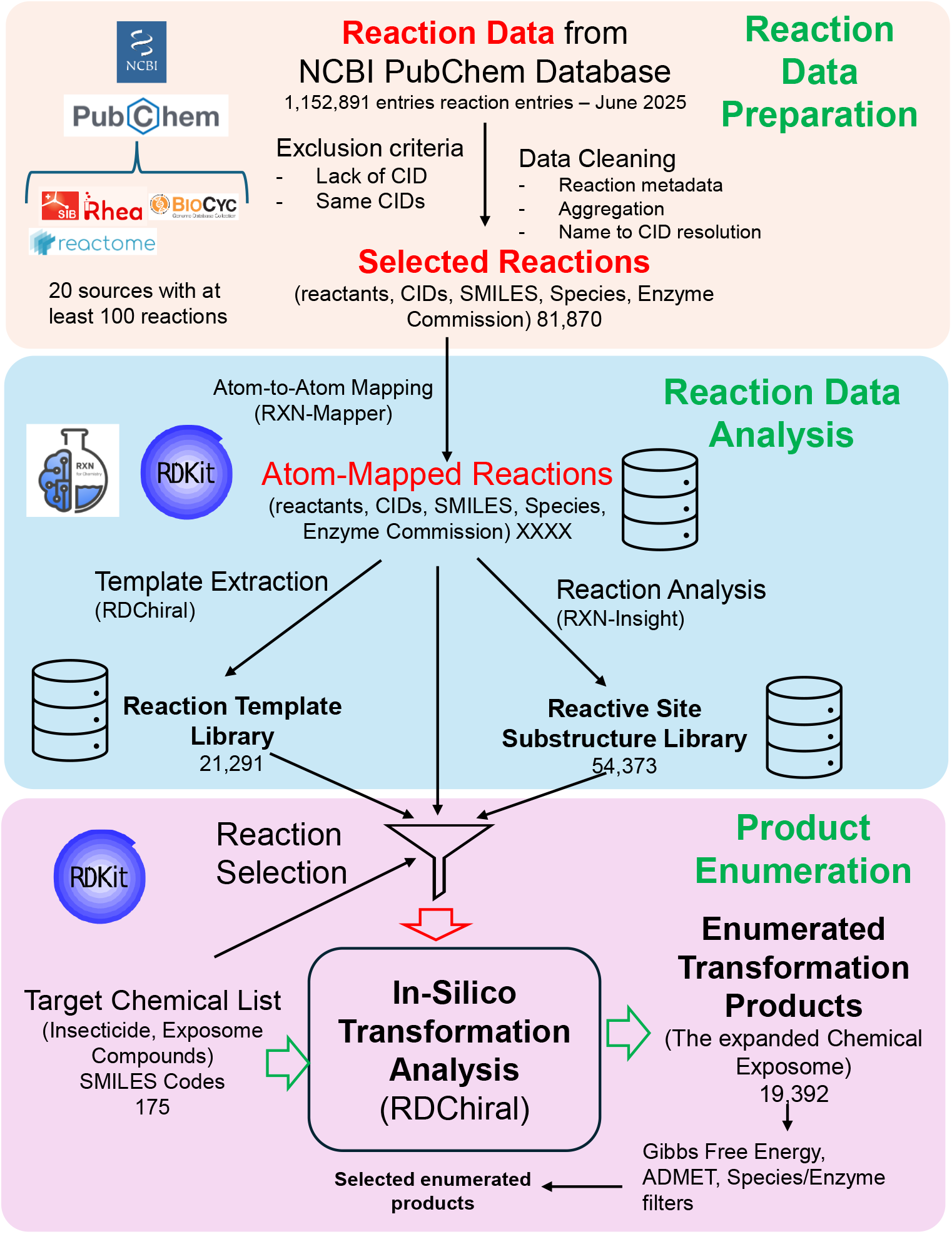
Overview of the In-silico transformation analysis (ISTA) workflow. Reaction data is aggregated from the NCBI PubChem database, incorporating sources such as Rhea, BioCyc, and Reactome. The raw data undergoes cleaning and exclusion criteria (e.g., removing entries with missing or duplicate CIDs) to yield a curated set of selected reactions. The selected reactions are then processed using RXNMapper to generate atom-to-atom mappings. This mapped data is then analyzed to create two key libraries: a Reaction Template Library generated via RDChiral (containing 21,291 templates) and a Reactive Site Substructure Library generated via Rxn-INSIGHT (containing 54,373 substructures). Next, the generated libraries are applied to a target list of chemicals (e.g., insecticides) to predict potential biotransformations. The In-Silico Transformation Analysis enumerates 19,392 unique transformation products, which are subsequently prioritized using filters for Gibbs Free Energy, ADMET properties, and species/enzyme associations.

## Materials and Methods

### Reaction Data Preparation

Reaction data were sourced from the National Center for Biotechnology Information (NCBI) PubChem database (https://pubchem.ncbi.nlm.nih.gov/), a publicly available repository of chemical information, including structures, properties, and biological activities. Reactions were submitted by different sources including RHEA^22^, Reactome^23^, PharmaKGB^24^ and BioCyc^25^ along with biological organism annotations (Table S1 and S2). For each reaction, reactant and product fields contain chemical names and CIDs that were mapped to the reaction. Since, these were crowd-sourced reaction data, we passed it through two exclusion criteria. First, we removed all the reactions in which the counts of mapped CIDs and chemical names were not same, meaning one of the chemical names was not connected with a chemical structure. Second, we removed reactions in which same CID appeared on both sides, which could be an error by data submitted to the PubChem. Then, the canonical SMILES of all reactants and products in a reaction were concatenated with ‘.’ and then joined using the ‘>>’ separator to create a reaction SMILES (e.g. “SMILES 1>>SMILES 2”). To ensure each chemical transformation was represented only once, we obtained standardized compound identifiers (CIDs) for both reactants and products and combined them to create a unique reaction CID combination. This approach identified duplicate and redundant reactions that may occur when the same transformation is reported multiple times across different data sources, for example a reaction conserved across multiple species will have more than one entry. Full reaction data are available at Zenodo accession (https://zenodo.org/records/18530567, #Table 1)

### Chemical Reaction Analysis

To analyze chemical reactions particularly reaction classification, naming, capturing transformation patterns, we utilized the Rxn-INSIGHT^26^, a python library which uses reaction SMILES as input. We processed the filtered and non-duplicate reaction set from previous step through Rxn-INSIGHT library. Our goal was to obtain the atom-to-atom mapping (AAM) of reactions computed by the RXNMapper^20^, python library within the Rxn-INSIGHT workflow. Atomic mapping enables tracking of atoms from substrates to products within a reaction, allowing an accurate analysis of chemical transformation data. RXNMapper has been shown to outperform other AAM tools^27^. In an atomically mapped reaction, atom map numbers are assigned in a sequential manner in the context of reaction (i.e. based on how reaction data is constructed), from 1 to the maximum atoms in a reaction (Figure 2). To sanitize a reaction, Rxn-INSIGHT checks valency, aromaticity, kekulization, atom hybridization and formal charges for the individual SMILES within a reaction. The sanitized mapped reaction is considered for further analysis.

**Figure 2.**
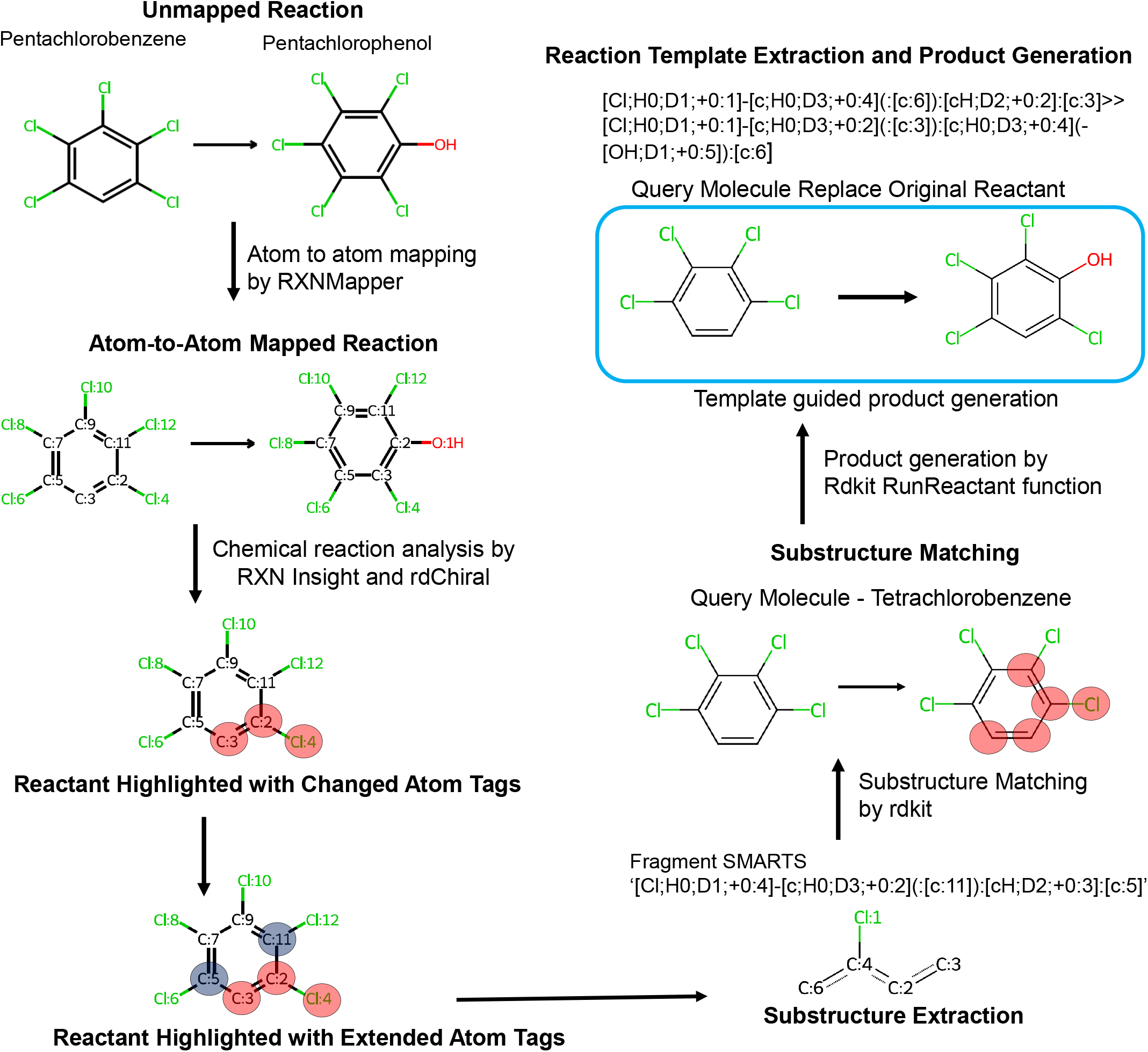
Template guided enumeration of transformation products. The workflow begins with atom-to-atom mapping of archived chemical/biochemical reactions using RXNMapper. From these mapped reactions, reactive sites, atoms that undergo changes during the reaction were identified, and corresponding substructures were extracted from the reactants. The chemical transformation templates that define transformation patterns were generated from atom-to-atom mapped reaction using the rdChiral, a python library. For a given query compound, substructure matching was performed using RDKit and corresponding transformation template was applied to generate potential transformation products. The workflow is exemplified by the transformation of pentachlorobenzene into pentachlorophenol.

Rxn-INSIGHT also detects the functional groups of individual reactants and products by matching molecular substructure of predefined list of most common functional groups (n=107) in SMARTS notation. In addition, Rxn-INSIGHT calculated several other characteristics for a reaction, including direct participation of ring reactants in transformation, scaffold, involvement of NOS (nitrogen, oxygen, and sulfur) in reaction center, number of reactants and products, and the identification of unaltered molecules such as reagents, solvents, catalysts that do not undergo structural changes during reaction. Each reaction was uniquely identified by an RXN_ID field and included its reaction representation in unmapped reaction (REACTION_SMILES) and atom-to-atom mapped reactions (MAPPED_REACTION, SANITIZED_MAPPED_REACTION). The output of Rxn-INSIGHT workflow is available in the Zenodo (https://zenodo.org/records/18530567, #Table 2).

### Reaction template and substructure database generation

To capture changes in the connectivity between products and their corresponding reactants, a reaction template was generated from each sanitized mapped reaction using the RDChiral library^21^. These templates indicate a chemical transformation pattern that describes how reactants converted to products during a chemical reaction. These templates were encoded in the SMARTS format, which captured key reactive substructures and facilitated generalization, allowing for application of these rules to a wide variety of compounds with similar or identical substructures. We extracted substructures representing chemical transformation sites within individual reactants for each sanitized mapped reaction using the RDChiral python library^21^. These substructures were needed to screen the reactions for a new substrate. Next, “get_changed_atoms”, a helper function from the RDChiral library was used to identify changed atoms by comparing local atomic properties such as atomic number, number of hydrogens, formal charge, degree, number of radical electrons, aromaticity, or the identity of neighboring atom-differ between reactants and products. To accurately capture the structural context of a compound’s reactivity, all atoms initially identified as changed atoms were expanded (radius=1) by including their directly connected neighboring atoms. Furthermore, if any of these atoms, whether originally changed or their direct neighbors were part of a predefined list of 30 known functional groups (e.g., alkenes, carbonyls, carboxylic acids), the entire functional group was included in the set of changed atoms. Functional groups were considered part of the reaction center if at least one of their atoms was involved in the set of transformed atoms. This expansion ensures that the chemical context influencing reactivity is preserved, as reactivity often arises from interactions among atoms within their immediate chemical environment. For every reaction, the substructure corresponding to individual reactant was generated as a SMARTS pattern. The substructure SMARTS was annotated with its corresponding SMILES representation, reaction ID, the originally identified changed atoms, and the list of expanded atoms derived from structural context analysis. The substructure and template databases are available at Zenodo (https://zenodo.org/records/18530567, #Table 3).

### In-silico transformation analysis for insecticides

A total of 181 insecticides (https://zenodo.org/records/18530567, #Table 4) were selected for template-based biotransformation product enumeration analysis. These compounds were chosen based on three primary criteria: chemical diversity, agricultural significance, and structural availability. Based on the structural properties, mode of entry, mode of action, and toxicity profiles, the selected insecticides were categorized into twelve different classes as follows: arsenical insecticides (n=2), carbamate insecticides (n=24), neonicotinoid insecticides (n=7), nereistoxin insecticides (n=3), organochlorine insecticides (n=6), organophosphorus insecticides (n=56), phosphoramido insecticides (n=4), pyrazole insecticides (n=5), pyrethroid insecticides (n=27), acaricides (n=24), nematicides(n=4) and others (n=19). Additionally, compounds with defined chemical structures and available canonical SMILES were considered for this study. SMILES notation representing chemical structures for selected insecticides was downloaded from PubChem database and validated using the RDKit library to ensure structural consistency and format compatibility.

To predict the transformation of a query molecule, we first perform a substructure match of query molecules against a substructure library containing chemical transformation sites. This screening process identified the reactions and reaction templates relevant to the specific substructure. Then each template was applied to the queried insecticide using the ‘RunReactants’ function in RDkit. Products with highest similarity to the reactant were prioritized as the most relevant transformation product of the query compound. These product structures were queried against the PubChem database to see if they were already known structures. For the product, we then computed Gibbs free energy using Joback group-contribution approximation method, a widely used for predicting thermodynamic properties when empirical data are unavailable. We used Thermo python library (https://github.com/CalebBell/thermo) for thermodynamic calculations. For each insecticide, all the products were sorted by Gibbs free energy to rank them by thermodynamic stability. ADME properties for the product structures were computed using the ADMET-AI library^28^. The ADMET-AI is a new tool to predict 41 different ADMET properties. It is trained on experimental ADMET and toxicity data from the Therapeutics Data Commons (https://tdcommons.ai/). ADMET-AI predictions have shown better performance in comparison to several other models^28^.

Python scripts for the workflows are available at https://github.com/idslme/in-silico-transformation and we have provided a detailed description for these scripts in the supplementary section.

Computer resources: All the calculations were conducted using a Linux server with 24 threads (Intel(R) Xeon(R) CPU E5-2643 v2 @ 3.50GHz), 128 GB RAM and 1 TB of Solid-State Drive. Python version Python 3.7 was used to develop the scripts. It took approximately 8 hours to complete the enumeration for 175 insecticides.

## Results

### Chemical Reaction Database and Analysis

We have assembled a comprehensive informatic workflow to enumerate transformation products for an exposome chemical by integrating publicly available reaction chemistry data and available cheminformatic methods (Figure 1). A total of 1,152,892 reactions were retrieved from the PubChem database in June 2025 (https://zenodo.org/records/18530567, #Table 1). There were many reactions that had more than one entry due to multiple species and databases covering the same reaction. After applying the exclusion criteria, 81,780 unique reactions were selected. 65% reactions were linked with Human (Taxonomy ID:9606), 17% were abiotic and rest were biological but without any specific taxonomy. 79.2% of reactions had an enzyme commission number annotation. Major reaction sources were PathBank, RHEA, BioCyc, PlantCyc, Reactome, MetXBioDB, WikiPathways, HSDB, EAWAG, INOH, and PhamGKB (https://zenodo.org/records/18530567, #Table 2 and Table S1). As expected, model organisms were the most frequently associated with a reaction (Table S2). Table S1 and S2 also shows the proportion of reactions that were excluded for major reaction sources and organisms due to missing mapping between a chemical name and PubChem CID. The Rxn-INSIGHT library was used to categorize each reaction into one of ten major classes. The most prevalent class was protection, followed by acylation and C-C coupling (Figure 3A). To learn about the unique reaction chemistries, we summarize the reaction templates by each reaction class. A reaction template captures and tracks the atomic changes between reactant and product (Figure 2). Using the RDChiral library^21^, we extracted the templates for each atom-mapped reaction. The strength of this template-based approach is that the predicted transformations are mechanistically interpretable. Within each reaction class, a majority of unique templates were obtained from enzymatic reactions (Figure 3B). Functional group analysis of both reactants and products revealed the presence of 90 functional groups in reactants and products, with secondary alcohols, ethers, and esters being the most observed in both (https://zenodo.org/records/18530567, #Table 2). 18.4% of reactants and 21% of products contain ring structures; 8.3% of reactions involve changes in ring structure; and 86.4% of molecules were detected with a Markov scaffold. A substantial portion of the templates were obtained from enzymatic (57%), with the remainder from non-enzymatic reactions (Figure 3C), indicating that our enumeration approach is biased toward biological reactions. We have over 80,000 reactions to test for each chemical, many of them would not yield products for a chemical. To facilitate the faster matching of a query compound, sub-structures that correspond to the metabolic hotspots or reaction centers were extracted for each reaction (See method and https://zenodo.org/records/18530567, #Table 3). We use this substructure library to screen for reactions that have the relevant reactive site in the query chemical (Figure 1 and 2).

**Figure 3.**
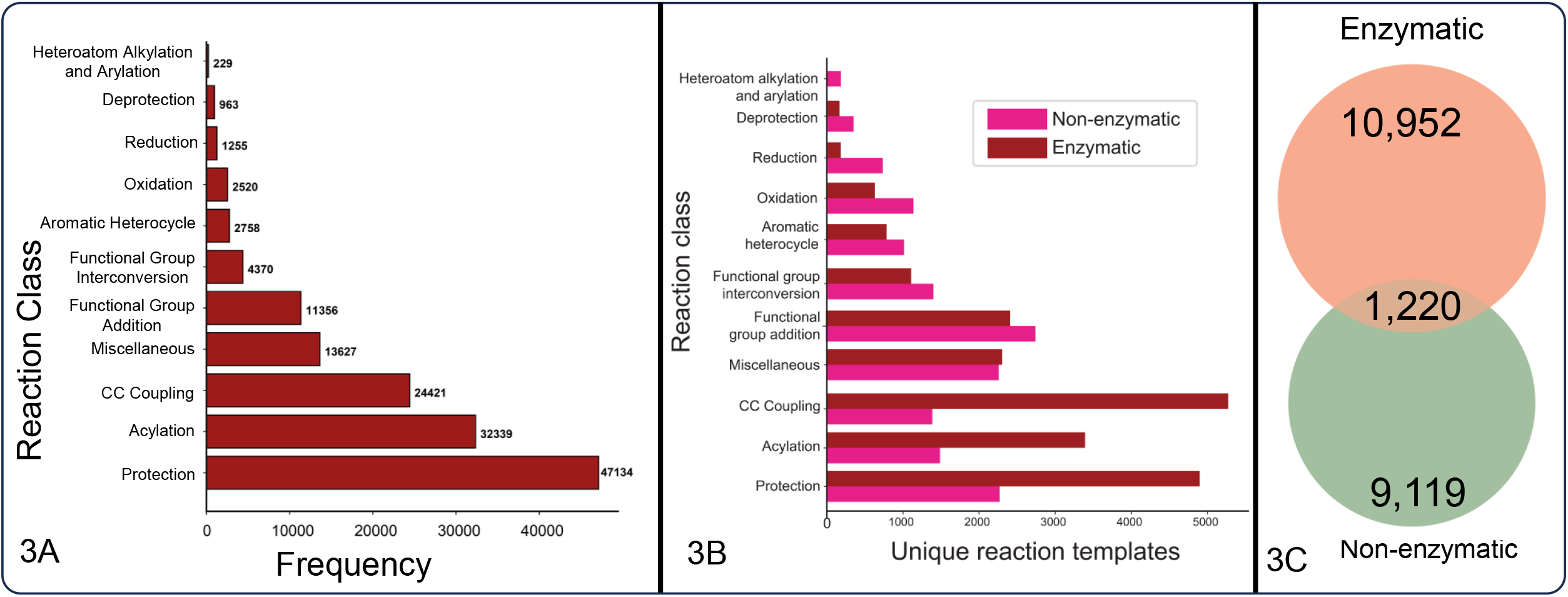
Overview of the reaction data sourced from the PubChem Database. Figure 2A shows the total number of reactions for each reaction class, for which transformation template has been generated. 3B shows the count of unique reaction templates. 3C represents distribution of unique chemical transformation templates extracted from enzymatic and non-enzymatic reactions, including the number of overlapping shared between the two categories.

### In-silico transformation analysis for insecticides

For the present study, 181 insecticides with a wide range of molecular characteristics reflecting significant physicochemical and structural diversity were chosen for template-guided in-silico transformation analysis (https://zenodo.org/records/18530567, #Table 4). These cover historically and currently significant insecticides like parathion, pentachlorophenol, carbaryl, rotenone, and dicofol. The chemical diversity of insecticides is illustrated in Figure S1. While a few subclasses exhibit high intra-class similarity, the majority are widely dispersed, indicating substantial structural divergence across the chemical space. Organophosphorus, carbamate, and pyrethroid insecticides formed distinct groups in a tSNE-based chemical diversity visualization using molecular fingerprints. There is some overlap between acaricides, others, carbamate, and phosphoramido insecticides, indicating certain overlapping structural characteristics. We expected to identify different chemical reactions that can transform these insecticides due to the chemical diversity.

The transformation product enumeration workflow (Figure 1 and 2) was applied to a structurally diverse set of 181 insecticides. The workflow successfully generated transformation products for 175 of the 181 query compounds. This analysis yielded a total of 19,392 unique transformation products (https://zenodo.org/records/18530567, Table 5 and Table 8) as the major product. Total products were 36,600 including side products, such as ethanol, water and other smaller side products. The number of major products enumerated for each insecticide varied significantly, with a median of 62 unique products per compound (Figure S2). Pyrethroid insecticides were particularly reactive in the model, contributing over 4,350 unique products as a class (Figure 4A), followed by Acaricides (4,051) and Organophosphorus (3,740) (Figure S2). Querying against the PubChem database revealed that 1,249 (6.4%) of the enumerated products were already known chemical structures, though many had not been previously linked to their parent insecticides. For example, Nicotine (PubChem CID 89594) has 60 (32%) of its enumerated transformation products corresponding to known compounds in PubChem. Overall, 209 (16.7%) unique PubChem CIDs are associated with at least one known reaction. Within this subset, a substantial fraction exhibited documented reactions with the parent compounds (Figure 4A and Figure S3).

**Figure 4.**
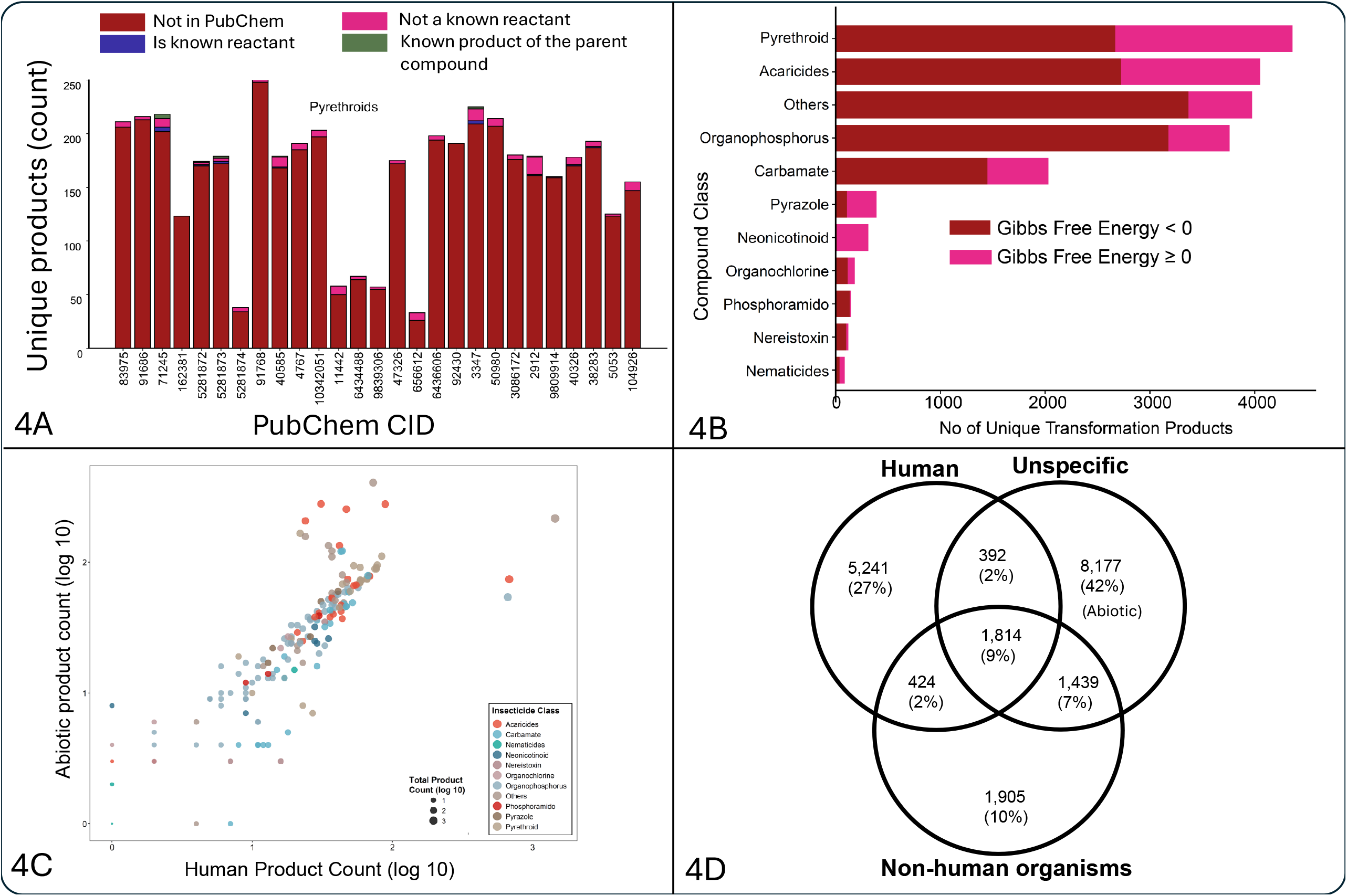
Analysis of Transformation Products of Pesticides. (4A) Unique products per compound. It shows the count of unique transformation products for selected parent compounds identified by their PubChem CID for Pyrethroids class. The colors indicate the status of each product: Not in PubChem (dark red), Not a known reactant (pink), Is a known reactant (dark blue), and Known product of the parent compound (green). (4B) Transformation products by compound class and thermodynamics. It displays the total number of unique transformation products categorized by compound class. It shows the proportion of reactions with a change in Gibbs Free Energy < 0 (spontaneous, dark red) and ≥ 0 (non-spontaneous, pink). (4C) Comparison of human and abiotic product counts. It compares the log10-transformed counts of products formed through human metabolism (x-axis) versus abiotic degradation (y-axis). The points are colored according to the Insecticide class, and their size is proportional to the log10-transformed total product count. (4D) Overlap of transformation products across systems. The Venn diagram illustrates the distribution of transformation products based on the systems in which they are formed: Human, Non-human organisms, and Unspecific (Abiotic). The diagram shows the number and percentage of products that are unique to each system or shared between them.

We checked the enumerated products for 20 insecticides against the predicted products by the BioTransformer^29^ online tool. BioTransformer generated 409 total, and 259 unique products for these selected 20 insecticides (Zenodo, Table 7). 229 (89%) products were covered by our enumerated product list that had both side products and major products. BioTransformer uses curated, specific rules, both enzymatic and abiotic to generate products for a query chemical. Our approach covers a majority of products predicted by BioTransformer, but we have additional enumerated products (SI Table 5) which provide hypotheses for discovering new reaction rules.

We also cross-validated the enumeration of transformation products by checking the literature in which the product chemicals have been previously reported. We could link back products of 65 insecticides and 136 products to published literature (Table S5). Known oxon metabolites including Chlorpyrifos oxon^30^, Malaoxon^31^, Paraoxon^32^, glucuronides, including Pentachlorophenol glucuronide^33^ and nicotine N-glucuronide^34^ were retained in our results, supporting that overall enumeration approach is useful and can be relevant for extending the chemical exposome. Enumerated products that are linked with PubChem CIDs (1,249) represents known and highly plausible chemical structures and reactions which have not been published yet, suggesting high-priority hypotheses.

### Characterization and Prioritization of Transformation Products

To assess the biological plausibility for the enumerated transformation products, we checked the reaction sources, thermodynamic stability and ADME properties. To rank the products, we have included Gibbs free energy calculations. More than half of the enumerated products were thermodynamically stable (Figure 4B). Insecticides from the Others and Organophosphorus categories showed highest proportion of products with Gibbs Free energy < 0, whereas Neonicotinoid has most products with Gibbs Free Energy ≥ 0. For each insecticide class, the ratio between human vs abiotic reaction products were similar, with some notable exception in Acaricides which has more abiotic products and few pyrethroids which had more human products (Figure 4C). Overall, up to 40% of enumerated products were linked with a Human, 42% with abiotic and rest were un-specific taxonomy reactions (Figure 4D). We also computed the ADME properties for the enumerated products using ADMET-AI^28^ library which predicted 41 endpoints for a chemical (Table S4). Many of the known toxic metabolites of insecticides such as oxons have overall ADME properties of a toxic chemical. Pyrethroids^35^ and organophosphates^36^, been shown to induce liver injury through oxidative stress and inflammation^37^, supporting a higher predicted drug-induced liver injury score (https://zenodo.org/records/18530567, Table 6). Overall, this combination of thermodynamic stability, biological sources, and ADME properties are helpful in prioritizing the enumerated compounds for future experimental and computational analyses.

## Discussion

This study presents a new transformation product enumeration workflow that utilizes >80,000 entries of PubChem reaction data to generate templates for in-silico transformation analysis. By applying this workflow to 181 insecticides, we predicted 19,392 unique major transformation products, substantially expanding the known chemical space for this important class of environmental exposures. Since we are using a very large number of reactions, the presented approach should be used for enumerating products for a given chemical. A key finding was that while 1,249 of these products were known compounds in PubChem, many had not been previously associated with their parent insecticides, representing new transformation hypotheses. Furthermore, the generation of thousands of entirely new chemical structures highlights the workflow’s potential in developing experimental and computational methods to illuminate the “dark matter” of the chemical exposome. The integration of thermodynamic calculations, species and enzyme commission information, ADME-Tox property prediction provides a prioritization approach, allowing researchers to refine this large list of potential products into a smaller, manageable list of high-priority candidates for further investigations.

To demonstrate the utility of the workflow, a chemically diverse set of 181 insecticides was selected to investigate their potential transformations. Insecticides are used widely in agricultural, domestic, and public health settings. While they are crucial for crop yield, there is growing concern over their effects on non-target organisms and the environment^38-40^. Human exposure occurs through various routes, including occupational handling and low-level chronic exposure from food, water, and household dust. Human biomonitoring studies have confirmed ubiquitous exposure to classes like organophosphates and pyrethroids by measuring their transformation products in urine^38, 41^. The enumerated transformation product database generated in this work can directly enhance these biomonitoring efforts by expanding targeted screening panels and improving the interpretation of untargeted datasets where transformation products often correlate with parent chemicals.

Our study has introduced the application of the reaction data from PubChem^19^ in creating a template library of reactions and applied to an important chemical class in the exposome (Figure 1). This extensive data set enabled the extraction of a diverse array of transformation templates, capturing a broad spectrum of known chemical and biological reactions. These templates facilitated the systematic enumeration of transformation products by matching the reactive sites of insecticides with known reaction patterns, allowing for comprehensive simulation of potential transformation products. Each predicted metabolite was subsequently queried against the PubChem database to determine whether it corresponded to a known compound or represented a novel, previously uncharacterized entity. To our knowledge, this is first time an enumeration approach of this scale using known reactions is created.

In-silico transformation may facilitate annotation of “chemical dark matter” in untargeted metabolomics and exposomics datasets by generating probable biotransformation products of parent compounds. Our proposed strategy to use thermodynamic stability, species, enzyme and ADME properties can narrow down thousands of possible structures to a manageable set. This may provide new hypothesis regarding the transformation of xenobiotics. These new structures can be used for formula matching, in-silico spectra prediction, and retention time prediction^42^ which can aid in the annotation of untargeted high resolution mass spectrometry datasets^43^.

We acknowledge that 94% of the enumerated transformation products for the tested insecticides were not yet found in the PubChem database, rendering them hypothetical structures. Biological plausibility of these structures needs further experimentation, however having the list of these hypothetical structures available may serve as a useful resource for the non-targeted screening of these transformation products in environmental and biological samples. Among the predicted products with CIDs, 136 of them have been reported in the literature, for example, 3-vinylpyridine^44^, paraoxon^44^, 3-chlorophenol^45^, 1-naphthol^46^ and . Retrieval of these known products supports the usefulness by our workflow for enumerating new hypotheses regarding transformation reactions of a compound from the chemical exposome.

The approach presented is limited to predicting reaction products and it cannot model enzyme kinetics or concentration-dependent effects of a substrate. Our approach cannot model 3D structures. Stereochemical changes alter spatial configuration without affecting atom connectivity, which limits the accuracy of 2D-based atom-to-atom mapping tools like RXNMapper. The current workflow supports screening of reactive sites and plausible transformation products. However, it lacks enzyme kinetics and thermodynamic constraints, limiting its ability to rank competing pathways.

The current workflow relies on the crowd-sourced reaction data available. We kept the workflow as general as possible for the broad coverage of chemical reactions that may expand the chemical exposome. By utilizing available species information and EC annotations, we can constrain the enumerated products by different context, such as products that are exclusive to Human biochemistry. However, there is a need to curate and collate, probably in a semi-automated way, all the reactions that have been deposited in the PubChem reaction database.

In comparison to drug metabolism studies, which focus on phase I and phase II transformations of a single chemical, we have taken a much broader approach to cover reactions from different databases that cover primary metabolism, abiotic, xenobiotic transformation. This allowed us to expand the chemical exposome, as shown by the test study for insecticides. We did rely on the cheminformatics tools (RXNMmapper, Rxn-INSIGHT and RDChiral) that have been developed mainly for studying drug metabolism, but we have first time applied these tools and the conceptual product enumerating workflow for a large set of chemicals that important for the chemical exposome.

The next critical step is to leverage these enumerations to query existing untargeted metabolomics data from human cohorts, animal models, or environmental samples. Finding evidence of these computationally predicted transformation products in real-world biospecimens would provide strong validation for the workflow and confirm the presence and significance of these new components of the chemical exposome. We have only tested for a set of insecticides. However, the proposed computational method to enumerate in-silico transformation products may be useful for other chemical classes in the exposome and follow up analyses for those chemicals will be useful for expanding existing exposomic databases.

## Conclusion

We have integrated existing reaction data from PubChem and cheminformatics software to develop a new workflow that can enumerate the transformation products for an exposome chemical. We have demonstrated the application of this in-silico transformation analysis (ISTA) workflow to expand the chemical space of insecticides. This workflow can expand the chemical exposome databases by many folds, enabling novel hypothesis regarding xenobiotic transformation, and will be helpful in better understanding the role of environmental toxins and chemicals in human health.

## Funding

The work was in part supported by U24ES035386, R24ES036917, R01ES035478, P30ES023515, UL1TR004419, R01ES032831 and R01ES033688.

## Conflict of interest

The authors declare no competing financial interest.

## Author’s contribution

MJ and DB planned the study, implemented the software, prepared the results and drafted the manuscript. All authors have reviewed the manuscript’s content.

## Supporting information

Additional tables and figures are available in the SI section

## Code availability

All in-house scripts are available at https://github.com/idslme/in-silico-transformation

## Data availability

Data are available at Zenodo https://zenodo.org/records/18530567

## Notes

https://zenodo.org/records/15881389

### Competing Interest Statement

The authors have declared no competing interest.

### Summary of Updates

The manuscript was revised in response to the reviewer's comment.

