## Supplementary material for "Enumerating the chemical exposome using in-silico transformation analysis – an example using insecticides": SI Tables and Figures

**Figure S1**: Visualization of Insecticides Chemical Space Using **Stochastic Neighbor Embedding (t-SNE).** The t-SNE projection of insecticides based molecular fingerprints shows structural and chemical diversity between and within the insecticide subclasses.

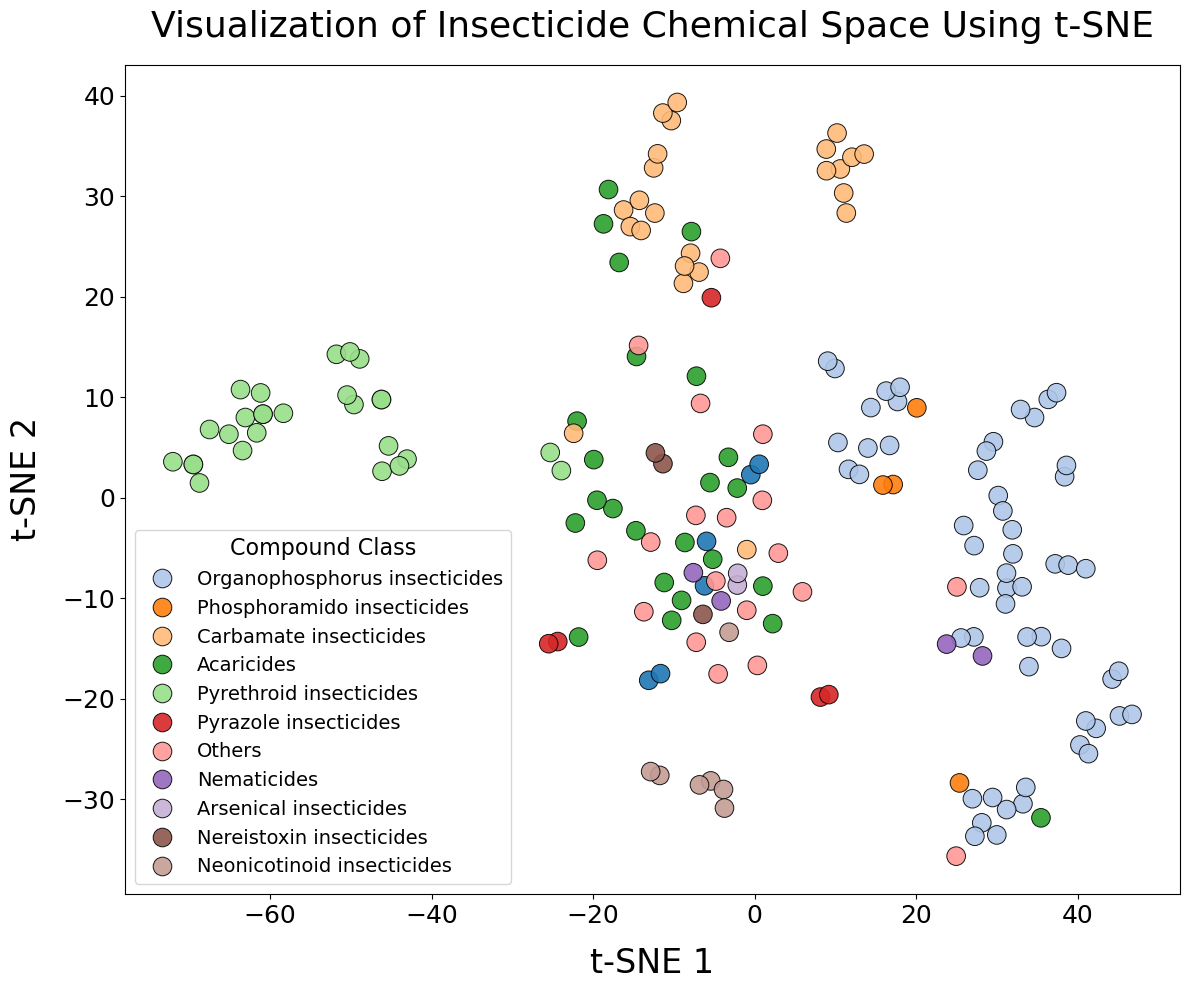

**Figure S2**: **Transformation Product Counts,** shows total number of unique transformation products generated for each query compound in the insecticide subclasses. Pyrethroid class is shown in Figure 4a. Arsenical insecticides did not generate any product. Each stacked bar graph categorizes the transformation products as compounds enlisted in PubChem, not enlisted in PubChem, associated with any reaction, and overlapping reaction count with parent compound and transformation products.

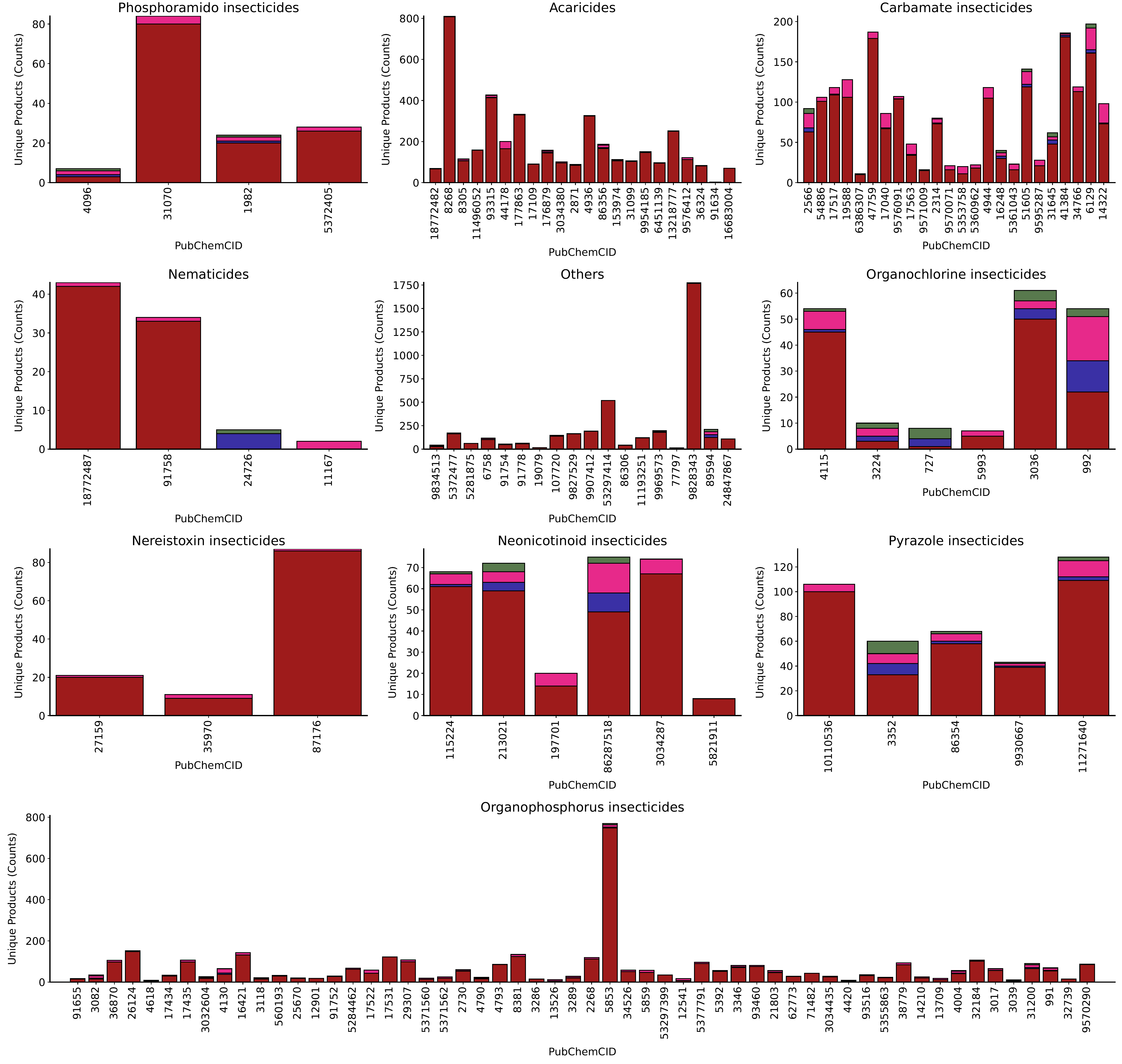

**SI Table 1**: **Reaction Sources and Distribution of Reaction Counts Across Sources,** summarized list of sources, including databases, publications or these from which each reaction entry was hosted in the PubChem database. For each source, it includes the total number of reactions, the number of unique reactions, and the subset of reactions known to occur in Homo sapiens. It also includes the number of unique Enzyme Commission (EC) numbers or enzyme names linked to each source to show reaction diversity. Source having at least 100 reactions are shown in this table.

| Source | Count | Unique Reactions by Source | Reaction in Homo Sapiens | Unique Reactions in Homo Sapiens | No of Unique EC Numbers | No of Reactions Missing Name to CID Map | Percentage Missing (%) |
| --- | --- | --- | --- | --- | --- | --- | --- |
| PathBank | 701922 | 85966 | 301973 | 57219 | 1593 | 157621 | 22.46 |
| PlantCyc | 396680 | 6206 |  |  | 943 | 118680 | 29.92 |
| Reactome | 205355 | 10922 | 28117 | 5339 | 1376 | 129297 | 62.96 |
| Plant Reactome | 146137 | 1542 |  |  | 125 | 40388 | 27.64 |
| BioCyc | 72665 | 13405 | 1831 | 1207 | 3328 | 17764 | 24.45 |
| RHEA | 17706 | 17423 |  |  | 6120 | 5595 | 31.6 |
| WikiPathways | 3974 | 3061 | 2761 | 2369 | 444 | 957 | 24.08 |
| MetXBioDB | 2126 | 2097 | 2099 | 2070 | 412 | 0 | 0 |
| COVID-19 Disease Map | 2046 | 1711 | 238 | 223 | 41 | 1116 | 54.55 |
| Environ. Sci. Technol. 58:22, 9828-9838 | 1606 | 1603 | 473 | 472 | 0 | 0 | 0 |
| HSDB | 1286 | 868 | 97 | 80 | 63 | 0 | 0 |
| Eawag-Soil | 1062 | 1062 |  |  | 0 | 0 | 0 |
| INOH | 972 | 612 | 792 | 557 | 506 | 189 | 19.44 |
| 10.5281/zenodo.4687924 | 294 | 294 |  |  | 0 | 0 | 0 |
| PharmGKB | 261 | 253 | 261 | 253 | 89 | 182 | 69.73 |
| Eawag | 204 | 204 |  |  | 0 | 0 | 0 |
| 10.1002/etc.3750 | 125 | 108 |  |  | 0 | 0 | 0 |
| 10.1021/acs.est.6b00140 | 123 | 105 |  |  | 0 | 0 | 0 |
| 10.1016/j.cbi.2005.06.007 | 108 | 108 |  |  | 7 | 0 | 0 |
| PPDB | 104 | 104 |  |  | 0 | 0 | 0 |

**SI Table 2**: **Distribution of Reaction Counts Across Biosystems,** shows the number of chemical and biochemical reactions associated with different biosystems, such as species and in vitro models. For each biosystem, we list the total number of reactions (Count), the number of unique reactions (No of Unique Reaction Count), and the data sources (Sources) that contribute to the reactions. Organisms with at least 1000 reactions are shown in this table.

| Biosystem | No of Reactions | Unique Reactions per Biosystem | Reaction Sources | No of Reactions Missing Name to CID Map | Percentage Missing (%) |
| --- | --- | --- | --- | --- | --- |
| Homo sapiens (human) | 335973 | 67075 | BioCyc\|COVID-19 Disease Map\|INOH\|PathBank\|PharmGKB\|Reactome\|WikiPathways | 38804 | 11.55 |
| Mus musculus (house mouse) | 98801 | 22773 | BioCyc\|INOH\|PathBank\|Reactome\|WikiPathways | 24551 | 24.85 |
| Rattus norvegicus (Norway rat) | 96459 | 21473 | PathBank\|Reactome\|WikiPathways | 23616 | 24.48 |
| Bos taurus (cattle) | 71823 | 17807 | PathBank\|WikiPathways | 10685 | 14.88 |
| Saccharomyces cerevisiae (baker's yeast) | 68029 | 12730 | PathBank\|WikiPathways | 51331 | 75.45 |
| Drosophila melanogaster (fruit fly) | 44929 | 9187 | BioCyc\|INOH\|PathBank\|Reactome | 27124 | 60.37 |
| Caenorhabditis elegans | 39238 | 8167 | INOH\|PathBank\|Reactome\|WikiPathways | 19178 | 48.88 |
| Arabidopsis thaliana (thale cress) | 25443 | 5957 | PathBank\|Plant Reactome\|WikiPathways | 8027 | 31.55 |
| Bos taurus (domestic cattle) | 18927 | 3735 | Reactome | 12308 | 65.03 |
| Sus scrofa (pig) | 18581 | 3653 | Reactome | 12071 | 64.96 |
| Canis lupus familiaris (dog) | 18074 | 3521 | Reactome\|WikiPathways | 11623 | 64.31 |
| Gallus gallus (chicken) | 16569 | 3238 | Reactome\|WikiPathways | 10461 | 63.14 |
| Xenopus tropicalis (tropical clawed frog) | 12507 | 2469 | Reactome | 7381 | 59.01 |
| Escherichia coli | 11528 | 2841 | PathBank\|WikiPathways | 8412 | 72.97 |
| Pseudomonas aeruginosa | 11369 | 3779 | HSDB\|PathBank | 7982 | 70.21 |
| Danio rerio (zebrafish) | 10185 | 2073 | Reactome\|WikiPathways | 5704 | 56 |
| Dictyostelium discoideum | 7252 | 1469 | Reactome | 3609 | 49.77 |
| Saccharomyces cerevisiae (brewer's yeast) | 5206 | 1060 | Reactome | 2859 | 54.92 |
| Solanum lycopersicum (tomato) | 5159 | 2759 | Plant Reactome\|PlantCyc | 1519 | 29.44 |
| Triticum aestivum (bread wheat) | 5156 | 2728 | Plant Reactome\|PlantCyc | 1534 | 29.75 |
| Schizosaccharomyces pombe (fission yeast) | 5134 | 1037 | Reactome | 2986 | 58.16 |
| Glycine max (soybean) | 5037 | 2560 | Plant Reactome\|PlantCyc | 1559 | 30.95 |
| Brassica napus (rape) | 5025 | 2682 | Plant Reactome\|PlantCyc | 1445 | 28.76 |
| Medicago truncatula (barrel medic) | 4999 | 2689 | Plant Reactome\|PlantCyc | 1406 | 28.13 |
| Aegilops tauschii | 4944 | 2591 | Plant Reactome\|PlantCyc | 1481 | 29.96 |
| Vitis vinifera (wine grape) | 4935 | 2617 | Plant Reactome\|PlantCyc | 1386 | 28.09 |
| Solanum tuberosum (potato) | 4932 | 2652 | Plant Reactome\|PlantCyc | 1473 | 29.87 |
| Populus trichocarpa (black cottonwood) | 4910 | 2623 | Plant Reactome\|PlantCyc\|WikiPathways | 1400 | 28.51 |
| Arabidopsis halleri | 4895 | 2635 | Plant Reactome\|PlantCyc | 1395 | 28.5 |
| Capsicum annuum | 4887 | 2623 | Plant Reactome\|PlantCyc | 1460 | 29.88 |
| Brassica rapa (field mustard) | 4874 | 2612 | Plant Reactome\|PlantCyc | 1406 | 28.85 |
| Cicer arietinum (chickpea) | 4857 | 2518 | Plant Reactome\|PlantCyc | 1366 | 28.12 |
| Panicum hallii | 4856 | 2525 | Plant Reactome\|PlantCyc | 1418 | 29.2 |
| Manihot esculenta (cassava) | 4829 | 2544 | Plant Reactome\|PlantCyc | 1361 | 28.18 |
| Sorghum bicolor (sorghum) | 4821 | 2473 | Plant Reactome\|PlantCyc | 1455 | 30.18 |
| Theobroma cacao (cacao) | 4816 | 2534 | Plant Reactome\|PlantCyc | 1418 | 29.44 |
| Citrus sinensis (sweet orange) | 4801 | 2514 | Plant Reactome\|PlantCyc | 1379 | 28.72 |
| Setaria italica (foxtail millet) | 4800 | 2477 | Plant Reactome\|PlantCyc | 1436 | 29.92 |
| Malus domestica (apple) | 4784 | 2579 | Plant Reactome\|PlantCyc | 1424 | 29.77 |
| Brachypodium distachyon (stiff brome) | 4779 | 2476 | Plant Reactome\|PlantCyc | 1367 | 28.6 |
| Prunus persica (peach) | 4776 | 2517 | Plant Reactome\|PlantCyc | 1403 | 29.38 |
| Gossypium raimondii (Peruvian cotton) | 4771 | 2483 | Plant Reactome\|PlantCyc | 1370 | 28.72 |
| Oryza glaberrima (African rice) | 4761 | 2448 | Plant Reactome\|PlantCyc | 1494 | 31.38 |
| Oryza brachyantha (malo sina) | 4751 | 2455 | Plant Reactome\|PlantCyc | 1408 | 29.64 |
| Helianthus annuus (common sunflower) | 4726 | 2502 | Plant Reactome\|PlantCyc | 1379 | 29.18 |
| Leersia perrieri | 4722 | 2435 | Plant Reactome\|PlantCyc | 1429 | 30.26 |
| Oryza punctata | 4709 | 2422 | Plant Reactome\|PlantCyc | 1411 | 29.96 |
| Oryza nivara | 4671 | 2380 | Plant Reactome\|PlantCyc | 1425 | 30.51 |
| Cucumis sativus (cucumber) | 4662 | 2431 | Plant Reactome\|PlantCyc | 1328 | 28.49 |
| Amborella trichopoda | 4642 | 2436 | Plant Reactome\|PlantCyc | 1370 | 29.51 |
| Cajanus cajan (pigeon pea) | 4633 | 2411 | Plant Reactome\|PlantCyc | 1368 | 29.53 |
| Musa acuminata (dwarf banana) | 4591 | 2405 | Plant Reactome\|PlantCyc | 1332 | 29.01 |
| Triticum urartu | 4567 | 2444 | Plant Reactome\|PlantCyc | 1338 | 29.3 |
| Brassica oleracea (wild cabbage) | 4520 | 2349 | Plant Reactome\|PlantCyc | 1351 | 29.89 |
| Fragaria vesca (wild strawberry) | 4463 | 2333 | Plant Reactome\|PlantCyc | 1292 | 28.95 |
| Erythranthe guttata (spotted monkey flower) | 4420 | 2358 | Plant Reactome\|PlantCyc | 1305 | 29.52 |
| Phoenix dactylifera (date palm) | 4403 | 2259 | Plant Reactome\|PlantCyc | 1263 | 28.68 |
| Dioscorea cayenensis subsp. rotundata (Guinea yam) | 4341 | 2368 | Plant Reactome\|PlantCyc | 1261 | 29.05 |
| Daucus carota (carrot) | 4310 | 2237 | Plant Reactome\|PlantCyc | 1250 | 29 |
| Selaginella moellendorffii | 4058 | 2050 | Plant Reactome\|PlantCyc | 1155 | 28.46 |
| Physcomitrium patens | 3986 | 1980 | Plant Reactome\|PlantCyc | 1041 | 26.12 |
| Oryza sativa (Asian cultivated rice) | 3812 | 1061 | Plant Reactome | 1210 | 31.74 |
| Nicotiana tabacum (common tobacco) | 3281 | 2218 | PlantCyc | 1036 | 31.58 |
| Chlamydomonas reinhardtii | 3173 | 1539 | Plant Reactome\|PlantCyc | 748 | 23.57 |
| Trifolium pratense (rotklee) | 3135 | 2161 | PlantCyc | 990 | 31.58 |
| Petunia axillaris (white moon petunia) | 3118 | 2131 | PlantCyc | 968 | 31.05 |
| Ricinus communis (castor bean) | 3108 | 2088 | PlantCyc | 926 | 29.79 |
| Zea mays subsp. mays (maize) | 3101 | 2013 | PlantCyc | 836 | 26.96 |
| Solanum pennellii | 3078 | 2081 | PlantCyc | 939 | 30.51 |
| Lotus japonicus | 3070 | 2104 | PlantCyc | 871 | 28.37 |
| Catharanthus roseus (Madagascar periwinkle) | 3037 | 2071 | PlantCyc | 894 | 29.44 |
| Brassica rapa subsp. pekinensis (Chinese cabbage) | 3028 | 2100 | PlantCyc | 865 | 28.57 |
| Oryza sativa Japonica Group (Japanese rice) | 3012 | 1950 | PlantCyc | 1051 | 34.89 |
| Arabidopsis lyrata (lyrate rockcress) | 3009 | 2067 | PlantCyc | 940 | 31.24 |
| Rosa multiflora (Japanese rose) | 3005 | 2086 | PlantCyc | 936 | 31.15 |
| Linum usitatissimum (flax) | 3003 | 2052 | PlantCyc | 909 | 30.27 |
| Camptotheca acuminata | 3001 | 2058 | PlantCyc | 835 | 27.82 |
| Rosa chinensis (China rose) | 2987 | 2058 | PlantCyc | 885 | 29.63 |
| Schrenkiella parvula | 2982 | 2041 | PlantCyc | 950 | 31.86 |
| Panicum virgatum (switchgrass) | 2967 | 2037 | PlantCyc | 909 | 30.64 |
| Miscanthus sinensis (eulalia) | 2964 | 2024 | PlantCyc | 896 | 30.23 |
| Spinacia oleracea (spinach) | 2953 | 1972 | PlantCyc | 855 | 28.95 |
| Boechera stricta | 2950 | 2033 | PlantCyc | 884 | 29.97 |
| Aquilegia coerulea (Rocky Mountain columbine) | 2941 | 1989 | PlantCyc | 886 | 30.13 |
| Brassica oleracea var. capitata (cabbage) | 2939 | 2045 | PlantCyc | 885 | 30.11 |
| Eucalyptus grandis (rose gum) | 2934 | 2041 | PlantCyc | 899 | 30.64 |
| Cannabis sativa | 2930 | 1981 | PlantCyc | 889 | 30.34 |
| Capsella rubella | 2926 | 2033 | PlantCyc | 893 | 30.52 |
| Eutrema salsugineum (saltwater cress) | 2924 | 2033 | PlantCyc | 893 | 30.54 |
| Anacardium occidentale (cashew) | 2916 | 1977 | PlantCyc | 899 | 30.83 |
| Capsella grandiflora | 2908 | 2008 | PlantCyc | 873 | 30.02 |
| Citrus x clementina (clementine) | 2907 | 1987 | PlantCyc | 858 | 29.51 |
| Salvia miltiorrhiza (redroot sage) | 2903 | 2031 | PlantCyc | 903 | 31.11 |
| Carica papaya (papaya) | 2901 | 2026 | PlantCyc | 929 | 32.02 |
| Phaseolus vulgaris (string bean) | 2895 | 1989 | PlantCyc | 910 | 31.43 |
| Oryza rufipogon (red rice) | 2887 | 1977 | PlantCyc | 894 | 30.97 |
| Saccharum spontaneum (wild sugarcane) | 2881 | 1942 | PlantCyc | 875 | 30.37 |
| Hevea brasiliensis (rubber tree) | 2880 | 1977 | PlantCyc | 896 | 31.11 |
| Beta vulgaris subsp. vulgaris (table beet) | 2877 | 1964 | PlantCyc | 877 | 30.48 |
| Chenopodium quinoa (quinoa) | 2870 | 1970 | PlantCyc | 890 | 31.01 |
| Setaria viridis | 2863 | 1936 | PlantCyc | 876 | 30.6 |
| Calotropis gigantea (mudar) | 2844 | 1962 | PlantCyc | 879 | 30.91 |
| Quercus lobata (valley oak) | 2844 | 1950 | PlantCyc | 850 | 29.89 |
| Dianthus caryophyllus (clove pink) | 2840 | 1949 | PlantCyc | 896 | 31.55 |
| Kalanchoe fedtschenkoi (South American air plant) | 2835 | 1937 | PlantCyc | 906 | 31.96 |
| Hordeum vulgare (barley) | 2810 | 1920 | PlantCyc | 819 | 29.15 |
| Quercus suber (cork oak) | 2804 | 1898 | PlantCyc | 844 | 30.1 |
| Corchorus capsularis (jute) | 2780 | 1873 | PlantCyc | 844 | 30.36 |
| Kalanchoe laxiflora | 2779 | 1920 | PlantCyc | 872 | 31.38 |
| Amaranthus hypochondriacus (grain amaranth) | 2743 | 1863 | PlantCyc | 844 | 30.77 |
| Ananas comosus (pineapple) | 2734 | 1851 | PlantCyc | 858 | 31.38 |
| Pyrus communis (pear) | 2725 | 1890 | PlantCyc | 786 | 28.84 |
| Humulus lupulus (European hop) | 2715 | 1852 | PlantCyc | 829 | 30.53 |
| Oryza longistaminata (red rice) | 2712 | 1838 | PlantCyc | 876 | 32.3 |
| Plasmodium falciparum (malaria parasite P. falciparum) | 2702 | 575 | Reactome\|WikiPathways | 1387 | 51.33 |
| Oryza barthii (African wild rice) | 2700 | 1849 | PlantCyc | 830 | 30.74 |
| Camellia sinensis var. sinensis | 2699 | 1851 | PlantCyc | 780 | 28.9 |
| Asparagus officinalis (garden asparagus) | 2696 | 1850 | PlantCyc | 838 | 31.08 |
| Prunus avium (sweet cherry) | 2674 | 1834 | PlantCyc | 855 | 31.97 |
| Lactuca sativa (garden lettuce) | 2664 | 1813 | PlantCyc | 832 | 31.23 |
| Human | 2664 | 2616 | 10.1021/acs.analchem.1c00142\|10.1021/acs.est.2c09854\|HSDB\|MetXBioDB\|Meyer C, Stravs MA, Hollender J (2024) Environ. Sci. Technol. 58:22, 9828-9838 | 0 | 0 |
| Phyllostachys edulis (mosochiku) | 2660 | 1788 | PlantCyc | 795 | 29.89 |
| Thalassiosira pseudonana CCMP1335 | 2654 | 1595 | BioCyc | 878 | 33.08 |
| Cuscuta campestris (field dodder) | 2650 | 1784 | PlantCyc | 776 | 29.28 |
| Spirodela polyrhiza (great duckweed) | 2639 | 1768 | PlantCyc | 789 | 29.9 |
| Oropetium thomaeum | 2630 | 1793 | PlantCyc | 797 | 30.3 |
| Tarenaya hassleriana (spider flower) | 2604 | 1773 | PlantCyc | 824 | 31.64 |
| Olea europaea (common olive) | 2587 | 1758 | PlantCyc | 800 | 30.92 |
| Musa balbisiana (Balbis banana) | 2565 | 1732 | PlantCyc | 788 | 30.72 |
| Cucumis melo (muskmelon) | 2565 | 1733 | PlantCyc | 777 | 30.29 |
| Senna tora | 2562 | 1749 | PlantCyc | 739 | 28.84 |
| Nelumbo nucifera (sacred lotus) | 2557 | 1737 | PlantCyc | 783 | 30.62 |
| Zostera marina | 2550 | 1702 | PlantCyc | 721 | 28.27 |
| Oryza meridionalis (Australian wild rice) | 2545 | 1759 | PlantCyc | 813 | 31.94 |
| Elaeis guineensis (African oil palm) | 2486 | 1684 | PlantCyc | 713 | 28.68 |
| Ostreococcus sp. 'lucimarinus' | 2406 | 1205 | Plant Reactome\|PlantCyc | 592 | 24.61 |
| Ginkgo biloba (maidenhair tree) | 2393 | 1609 | PlantCyc | 722 | 30.17 |
| Anthoceros agrestis | 2335 | 887 | PlantCyc | 543 | 23.25 |
| Sphagnum fallax | 2265 | 1481 | PlantCyc | 607 | 26.8 |
| Streptomyces coelicolor A3(2) | 2182 | 1028 | BioCyc | 389 | 17.83 |
| Marchantia polymorpha (liverwort) | 2101 | 1385 | PlantCyc | 579 | 27.56 |
| Oryza sativa Indica Group (long-grained rice) | 2008 | 553 | Plant Reactome | 590 | 29.38 |
| Citrullus lanatus (watermelon) | 1999 | 1340 | PlantCyc | 488 | 24.41 |
| Panicum hallii var. hallii | 1956 | 536 | Plant Reactome | 565 | 28.89 |
| Oryza barthii | 1954 | 539 | Plant Reactome | 545 | 27.89 |
| Oryza glumipatula | 1953 | 540 | Plant Reactome | 555 | 28.42 |
| Triticum dicoccoides | 1952 | 538 | Plant Reactome | 565 | 28.94 |
| Arachis duranensis | 1941 | 531 | Plant Reactome | 539 | 27.77 |
| Oryza rufipogon | 1939 | 532 | Plant Reactome | 577 | 29.76 |
| Zea mays | 1905 | 520 | Plant Reactome | 533 | 27.98 |
| Phaseolus vulgaris | 1876 | 521 | Plant Reactome | 512 | 27.29 |
| Triticum turgidum | 1871 | 516 | Plant Reactome | 529 | 28.27 |
| Lupinus angustifolius (narrow-leaved blue lupine) | 1862 | 516 | Plant Reactome | 510 | 27.39 |
| Arachis ipaensis | 1856 | 506 | Plant Reactome | 508 | 27.37 |
| Oryza longistaminata | 1854 | 510 | Plant Reactome | 520 | 28.05 |
| Actinidia chinensis | 1852 | 515 | Plant Reactome | 503 | 27.16 |
| Trifolium pratense | 1842 | 511 | Plant Reactome | 503 | 27.31 |
| Oryza minuta | 1841 | 504 | Plant Reactome | 517 | 28.08 |
| Coffea canephora | 1836 | 510 | Plant Reactome | 502 | 27.34 |
| Oryza officinalis | 1834 | 503 | Plant Reactome | 493 | 26.88 |
| Hordeum vulgare | 1830 | 505 | Plant Reactome | 507 | 27.7 |
| Arabidopsis lyrata | 1820 | 509 | Plant Reactome | 483 | 26.54 |
| Vigna angularis (adzuki bean) | 1815 | 506 | Plant Reactome | 494 | 27.22 |
| Severe acute respiratory syndrome coronavirus 2 | 1808 | 1488 | COVID-19 Disease Map | 995 | 55.03 |
| Beta vulgaris | 1801 | 499 | Plant Reactome | 500 | 27.76 |
| Oryza meyeriana var. granulata | 1789 | 491 | Plant Reactome | 490 | 27.39 |
| Oryza meridionalis | 1786 | 494 | Plant Reactome | 517 | 28.95 |
| Chromochloris zofingiensis | 1786 | 1186 | PlantCyc | 516 | 28.89 |
| Sequoia sempervirens (redwood) | 1776 | 1208 | PlantCyc | 479 | 26.97 |
| Volvox carteri | 1771 | 1169 | PlantCyc | 487 | 27.5 |
| Artemisia annua (sweet wormwood) | 1750 | 1215 | PlantCyc | 458 | 26.17 |
| Camelina sativa (false flax) | 1738 | 1234 | PlantCyc | 473 | 27.22 |
| Corchorus capsularis | 1730 | 479 | Plant Reactome | 465 | 26.88 |
| Oryza australiensis | 1709 | 472 | Plant Reactome | 473 | 27.68 |
| Vigna radiata | 1705 | 478 | Plant Reactome | 469 | 27.51 |
| Escherichia coli str. K-12 substr. MG1655 | 1703 | 853 | BioCyc | 268 | 15.74 |
| Eragrostis tef (tef) | 1693 | 1165 | PlantCyc | 427 | 25.22 |
| Escherichia coli CFT073 | 1686 | 793 | BioCyc | 444 | 26.33 |
| Picea abies (Norway spruce) | 1667 | 460 | Plant Reactome | 485 | 29.09 |
| Clostridium saccharoperbutylacetonicum N1-4(HMT) | 1656 | 873 | BioCyc | 476 | 28.74 |
| Coccomyxa subellipsoidea C-169 | 1633 | 1077 | PlantCyc | 445 | 27.25 |
| Chlorella variabilis | 1628 | 1042 | PlantCyc | 385 | 23.65 |
| Eucalyptus grandis | 1616 | 445 | Plant Reactome | 438 | 27.1 |
| Shigella flexneri 2a str. 2457T | 1591 | 721 | BioCyc | 231 | 14.52 |
| Salvia rosmarinus (rosemary) | 1591 | 1064 | PlantCyc | 408 | 25.64 |
| Escherichia coli O157:H7 str. EDL933 | 1590 | 752 | BioCyc | 432 | 27.17 |
| Jatropha curcas | 1582 | 438 | Plant Reactome | 424 | 26.8 |
| Micromonas commoda | 1556 | 1016 | PlantCyc | 404 | 25.96 |
| Nicotiana attenuata | 1552 | 431 | Plant Reactome | 428 | 27.58 |
| Escherichia coli B str. REL606 | 1549 | 776 | BioCyc | 166 | 10.72 |
| Allium sativum (garlic) | 1529 | 1035 | PlantCyc | 374 | 24.46 |
| Secale cereale (rye) | 1520 | 1051 | PlantCyc | 404 | 26.58 |
| Escherichia coli str. K-12 substr. W3110 | 1499 | 769 | BioCyc | 161 | 10.74 |
| Faidherbia albida | 1499 | 1050 | PlantCyc | 434 | 28.95 |
| Pinus taeda (loblolly pine) | 1491 | 408 | Plant Reactome | 431 | 28.91 |
| Artocarpus heterophyllus (jackfruit) | 1477 | 1029 | PlantCyc | 372 | 25.19 |
| Vigna subterranea (hog-peanut) | 1473 | 1037 | PlantCyc | 376 | 25.53 |
| Piper nigrum | 1454 | 1022 | PlantCyc | 398 | 27.37 |
| Micromonas pusilla | 1448 | 945 | PlantCyc | 368 | 25.41 |
| Artocarpus altilis (breadfruit) | 1427 | 983 | PlantCyc | 396 | 27.75 |
| Fragaria x ananassa (strawberry) | 1413 | 1009 | PlantCyc | 365 | 25.83 |
| Bacillus anthracis str. Ames | 1412 | 645 | BioCyc | 242 | 17.14 |
| Trypanosoma brucei | 1408 | 659 | BioCyc | 234 | 16.62 |
| Plasmodium falciparum 3D7 | 1406 | 656 | BioCyc | 353 | 25.11 |
| Mycobacterium tuberculosis CDC1551 | 1403 | 760 | BioCyc | 251 | 17.89 |
| Penicillium rubens Wisconsin 54-1255 | 1390 | 849 | BioCyc | 398 | 28.63 |
| Vibrio cholerae O1 biovar El Tor str. N16961 | 1383 | 649 | BioCyc | 234 | 16.92 |
| Lobularia maritima (sweet alyssum) | 1360 | 949 | PlantCyc | 373 | 27.43 |
| Aurantimonas manganoxydans SI85-9A1 | 1359 | 748 | BioCyc | 252 | 18.54 |
| Saccharomyces cerevisiae S288C | 1332 | 822 | BioCyc | 338 | 25.38 |
| Physalis pubescens (muyaca) | 1317 | 949 | PlantCyc | 351 | 26.65 |
| Persea americana var. drymifolia (Mexican avocado) | 1310 | 939 | PlantCyc | 331 | 25.27 |
| Draba nivalis | 1299 | 910 | PlantCyc | 269 | 20.71 |
| Caulobacter vibrioides NA1000 | 1295 | 797 | BioCyc | 1275 | 98.46 |
| Gnetum montanum | 1294 | 921 | PlantCyc | 334 | 25.81 |
| Nymphaea colorata (pocket water lily) | 1294 | 903 | PlantCyc | 331 | 25.58 |
| Caulobacter vibrioides CB15 | 1292 | 686 | BioCyc | 321 | 24.85 |
| Persea americana var. americana | 1290 | 879 | PlantCyc | 310 | 24.03 |
| Listeria monocytogenes 10403S | 1283 | 673 | BioCyc | 371 | 28.92 |
| Moringa oleifera (horseradish tree) | 1263 | 899 | PlantCyc | 326 | 25.81 |
| Bacteroides thetaiotaomicron VPI-5482 | 1257 | 686 | BioCyc | 229 | 18.22 |
| Vanilla planifolia (cultivated vanilla) | 1221 | 840 | PlantCyc | 292 | 23.91 |
| Plasmodium berghei yoelii | 1213 | 787 | PlantCyc | 226 | 18.63 |
| Anthoceros punctatus | 1185 | 806 | PlantCyc | 271 | 22.87 |
| Salvinia cucullata | 1175 | 789 | PlantCyc | 273 | 23.23 |
| Azolla filiculoides | 1171 | 794 | PlantCyc | 262 | 22.37 |
| Corynebacterium glutamicum ATCC 13032 | 1159 | 601 | BioCyc | 185 | 15.96 |
| Bacillus subtilis subsp. subtilis str. 168 | 1116 | 679 | BioCyc | 195 | 17.47 |
| Anthoceros angustus | 1110 | 770 | PlantCyc | 272 | 24.5 |
| Methylosinus trichosporium OB3b | 1055 | 638 | BioCyc | 271 | 25.69 |
| Synechococcus elongatus PCC 7942 = FACHB-805 | 1054 | 573 | BioCyc | 164 | 15.56 |

SI Table 3: **Transformation Product Generation Summary** represents an overview of PubChem CID of query compounds (Query CIDs) belonging to specific class of insecticides (Compound Class) and its transformation products data. It includes logarithm of the partition coefficient (logP) of query compounds (logP of Query CIDs) which indicates the compound’s hydrophobicity. The table shows the number of unique transformation products generated for each query compound, whether these products are already known in PubChem database (No of Transformation Product Metabolites Enlisted in PubChem), and the PubChem CID of transformation products if they are known (Transformation Product CIDs). It also lists CIDs that are linked to any reaction in the PubChem database (CIDs Associated with Any Reaction in PubChem) and CIDs that are in the PubChem database but not linked to any reaction (CIDs Not Associated with Any Reaction in PubChem). Additionally, it includes the reaction identifier for overlapping reactions between the query CID and transformation product CID (Overlapping Reaction ID Between Query CID and Transformation Product) and the total count of overlapping reactions between the query CID and transformation product (Overlapping Reaction Count Between Query CID and Transformation Product).

| Compound Name | Query CIDs | No of Transformation Product Metabolites Enlisted in Pubchem | Transformation Product CIDs | CIDs Associated with Any Reaction PubChem | CIDs Not Associated With Any Reaction PubChem | Overlapping Reaction ID Between Query CID and Transformation Product | Overlapping Reaction Count Between Query CID and Transformation Product | Compound Class |
| --- | --- | --- | --- | --- | --- | --- | --- | --- |
| Methamidophos | 4096 | 3 | [23413258, 57483867, 56605567] | [23413258] | [57483867, 56605567] | ['4096>>23413258'] | 1 | Phosphoramido |
| Cyenopyrafen | 18772482 | 2 | [141281160, 102401842] | [] | [141281160, 102401842] | [] | 0 | Acaricides |
| Carbofuran | 2566 | 23 | [129851137, 134096631, 15278, 20911, 134088367, 25210295, 154150841, 20923, 20924, 20925, 29374, 44098, 134109763, 27975, 76965064, 53662409, 76965065, 101200477, 27999, 24840801, 13591799, 15175289, 101708540] | [15278, 25210295, 27975, 27999, 15175289] | [129851137, 134096631, 20911, 134088367, 154150841, 20923, 20924, 20925, 29374, 44098, 134109763, 76965064, 53662409, 76965065, 101200477, 24840801, 13591799, 101708540] | ['1038\|2566\|962>>15278\|280\|644041', '2566>>15175289', '2566>>15278', '2566>>25210295', '2566>>27975', '2566>>27999'] | 6 | Carbamate |
| Imicyafos | 18772487 | 1 | [88095681] | [] | [88095681] | [] | 0 | Nematicides |
| Tetramethrin | 83975 | 5 | [78597, 21863245, 6455409, 2743, 20236475] | [] | [78597, 21863245, 6455409, 2743, 20236475] | [] | 0 | Pyrethroid |
| Chlorethoxyfos | 91655 | 2 | [73743736, 107199] | [] | [73743736, 107199] | [] | 0 | Organophosphorus |
| Dimethoate | 3082 | 15 | [14529280, 17345, 21721186, 118996707, 14209, 14210, 85772423, 14598891, 168140, 3080588, 89087503, 13553138, 3047253, 58808409, 199099] | [17345, 14210, 168140, 3080588] | [14529280, 21721186, 118996707, 14209, 85772423, 14598891, 89087503, 13553138, 3047253, 58808409, 199099] | ['17345>>3082', '3082>>14210', '3082>>168140', '3082>>3080588'] | 4 | Organophosphorus |
| Prothiofos | 36870 | 9 | [12894630, 20467943, 85933420, 186193, 12941237, 118710, 101614359, 12894620, 12894621] | [] | [12894630, 20467943, 85933420, 186193, 12941237, 118710, 101614359, 12894620, 12894621] | [] | 0 | Organophosphorus |
| Quinalphos | 26124 | 4 | [3014760, 101899713, 14526, 88258617] | [] | [3014760, 101899713, 14526, 88258617] | [] | 0 | Organophosphorus |
| Oxydemeton-methyl | 4618 | 4 | [17240, 157630968, 149432294, 157630967] | [] | [17240, 157630968, 149432294, 157630967] | [] | 0 | Organophosphorus |
| Flonicamid | 9834513 | 9 | [6923110, 76965321, 2777549, 86327757, 15766290, 2782643, 11651668, 2782930, 46835486] | [76965321, 2777549, 2782643, 2782930, 46835486] | [6923110, 86327757, 15766290, 11651668] | ['9834513>>2777549', '9834513>>2782643', '9834513>>2782930', '9834513>>46835486', '9834513>>76965321'] | 5 | Others |
| Methoxychlor | 4115 | 8 | [75048, 102399817, 92016457, 81869, 186510, 129714546, 86178806, 183679] | [183679] | [75048, 102399817, 92016457, 81869, 186510, 129714546, 86178806] | ['4115>>183679'] | 1 | Organochlorine |
| Cartap | 27159 | 1 | [153168834] | [] | [153168834] | [] | 0 | Nereistoxin |
| Thiacloprid | 115224 | 6 | [154824800, 132276193, 86222983, 87082633, 139792941, 9859386] | [86222983] | [154824800, 132276193, 87082633, 139792941, 9859386] | ['115224>>86222983'] | 1 | Neonicotinoid |
| Mecarbam | 17434 | 4 | [200728, 54328068, 16722134, 23618470] | [] | [200728, 54328068, 16722134, 23618470] | [] | 0 | Organophosphorus |
| Phenthoate | 17435 | 10 | [19398, 3732682, 106677356, 12630924, 4885809, 4885810, 12878836, 149436952, 10633916, 53680159] | [] | [19398, 3732682, 106677356, 12630924, 4885809, 4885810, 12878836, 149436952, 10633916, 53680159] | [] | 0 | Organophosphorus |
| Phosphamidon | 3032604 | 4 | [169493192, 3038082, 6444652, 16206879] | [169493192, 3038082, 6444652, 16206879] | [] | ['3032604>>16206879', '3032604>>169493192', '3032604>>3038082', '3032604>>6444652'] | 4 | Organophosphorus |
| Acetamiprid | 213021 | 9 | [102132322, 11094883, 102132323, 102132326, 11344811, 19967186, 73325014, 87081692, 59843549] | [11094883, 19967186, 73325014, 59843549] | [102132322, 102132323, 102132326, 11344811, 87081692] | ['213021>>11094883', '213021>>19967186', '213021>>59843549', '213021>>73325014'] | 4 | Neonicotinoid |
| Methyl Parathion | 4130 | 26 | [92291, 10372, 644235, 102206988, 13708, 17168, 4130, 121001, 100947629, 100947630, 85992250, 7485, 15859905, 85883589, 3039437, 3039439, 980, 86002008, 991, 31200, 190307, 86254309, 44152294, 173680, 13691762, 165631] | [644235, 4130, 7485, 980, 991, 31200] | [92291, 10372, 102206988, 13708, 17168, 121001, 100947629, 100947630, 85992250, 15859905, 85883589, 3039437, 3039439, 86002008, 190307, 86254309, 44152294, 173680, 13691762, 165631] | ['4130\|962>>1038\|3539116\|644235'] | 1 | Organophosphorus |
| Ethyl p-nitrophenyl benzenethiophosphonate | 16421 | 12 | [120450, 199491, 16421, 24841641, 24841642, 437293, 24841649, 53837970, 24841622, 23274938, 85701662, 67205183] | [] | [120450, 199491, 16421, 24841641, 24841642, 437293, 24841649, 53837970, 24841622, 23274938, 85701662, 67205183] | [] | 0 | Organophosphorus |
| Cycloprothrin | 91686 | 3 | [101614360, 101614361, 142991515] | [] | [101614360, 101614361, 142991515] | [] | 0 | Pyrethroid |
| Disulfoton | 3118 | 5 | [86109646, 54229683, 17241, 17242, 21666591] | [17242] | [86109646, 54229683, 17241, 21666591] | ['3118>>17242'] | 1 | Organophosphorus |
| Hydroprene | 5372477 | 9 | [12032832, 151372093, 44284358, 21522029, 44284182, 6437016, 22056699, 6374524, 152372445] | [] | [12032832, 151372093, 44284358, 21522029, 44284182, 6437016, 22056699, 6374524, 152372445] | [] | 0 | Others |
| Vamidothion | 560193 | 3 | [21123472, 54232686, 16212160] | [] | [21123472, 54232686, 16212160] | [] | 0 | Organophosphorus |
| Dinotefuran | 197701 | 6 | [71346722, 52952137, 16760148, 100958102, 87141530, 10932188] | [] | [71346722, 52952137, 16760148, 100958102, 87141530, 10932188] | [] | 0 | Neonicotinoid |
| Terbufos | 25670 | 2 | [25355, 41718] | [] | [25355, 41718] | [] | 0 | Organophosphorus |
| Tolfenpyrad | 10110536 | 6 | [15008516, 15008431, 123319859, 15008476, 15008478, 6966015] | [] | [15008516, 15008431, 123319859, 15008476, 15008478, 6966015] | [] | 0 | Pyrazole |
| Dicofol | 8268 | 3 | [145698, 8268, 160701] | [] | [145698, 8268, 160701] | [] | 0 | Acaricides |
| Etofenprox | 71245 | 12 | [23197441, 3067170, 3027267, 157700, 189485, 19972045, 19972046, 15590575, 19846640, 3067183, 153580853, 139596478] | [157700, 189485, 19972045, 139596478] | [23197441, 3067170, 3027267, 19972046, 15590575, 19846640, 3067183, 153580853] | ['71245>>139596478', '71245>>157700', '71245>>189485', '71245>>19972045'] | 4 | Pyrethroid |
| Bioresmethrin | 162381 | 0 | [] | [] | [] | [] | 0 | Pyrethroid |
| 2-Methylbiphenyl-3-ylmethyl (Z)-(1RS,3RS)-3-(2-chloro-3,3,3-trifluoroprop-1-enyl)-2,2-dimethylcyclopropanecarboxylate | 5281872 | 3 | [129865537, 596875, 167996382] | [129865537] | [596875, 167996382] | ['5281872>>129865537'] | 1 | Pyrethroid |
| Cyhalothrin | 5281873 | 5 | [139597315, 139594181, 13241320, 162599820, 89827927] | [139597315, 139594181] | [13241320, 162599820, 89827927] | ['5281873>>139594181', '5281873>>139597315'] | 2 | Pyrethroid |
| 2,3,5,6-tetrafluoro-4-methylbenzyl 3-[(1Z)-2-chloro-3,3,3-trifluoroprop-1-en-1-yl]-2,2-dimethylcyclopropanecarboxylate | 5281874 | 4 | [57490312, 6440522, 14404131, 12944026] | [] | [57490312, 6440522, 14404131, 12944026] | [] | 0 | Pyrethroid |
| Hydramethylnon | 5281875 | 1 | [15575087] | [] | [15575087] | [] | 0 | Others |
| Phosmet | 12901 | 0 | [] | [] | [] | [] | 0 | Organophosphorus |
| Benfuracarb | 54886 | 5 | [56991619, 148921192, 179117, 179118, 163808947] | [] | [56991619, 148921192, 179117, 179118, 163808947] | [] | 0 | Carbamate |
| Rotenone | 6758 | 14 | [442824, 70555849, 243725, 71176526, 92207, 182000, 100896946, 12315059, 139123155, 99189, 162977013, 68184, 9908730, 10431679] | [68184] | [442824, 70555849, 243725, 71176526, 92207, 182000, 100896946, 12315059, 139123155, 99189, 162977013, 9908730, 10431679] | ['6758>>68184'] | 1 | Others |
| Cadusafos | 91752 | 2 | [12518113, 145865100] | [] | [12518113, 145865100] | [] | 0 | Organophosphorus |
| Pyridaben | 91754 | 7 | [14348226, 14372649, 169451306, 14372650, 101091568, 53791569, 140984088] | [] | [14348226, 14372649, 169451306, 14372650, 101091568, 53791569, 140984088] | [] | 0 | Others |
| Isoprocarb | 17517 | 9 | [19842, 71353386, 54519087, 165346354, 42643, 12953311, 131849911, 23274041, 6943] | [6943] | [19842, 71353386, 54519087, 165346354, 42643, 12953311, 131849911, 23274041] | [] | 0 | Carbamate |
| Tetrachlorvinphos | 5284462 | 5 | [6538417, 6433329, 6913716, 20535960, 13032505] | [] | [6538417, 6433329, 6913716, 20535960, 13032505] | [] | 0 | Organophosphorus |
| Fosthiazate | 91758 | 1 | [11472025] | [] | [11472025] | [] | 0 | Nematicides |
| Tetradifon | 8305 | 9 | [695487, 7271, 277294, 22314677, 19017974, 14227574, 594650, 19802013, 19802015] | [7271] | [695487, 277294, 22314677, 19017974, 14227574, 594650, 19802013, 19802015] | [] | 0 | Acaricides |
| Cyanophos | 17522 | 15 | [6857921, 35771, 71373129, 23270156, 116078, 214704, 21128304, 70129, 85780053, 24841749, 3059191, 18788054, 43545, 13019, 85913022] | [] | [6857921, 35771, 71373129, 23270156, 116078, 214704, 21128304, 70129, 85780053, 24841749, 3059191, 18788054, 43545, 13019, 85913022] | [] | 0 | Organophosphorus |
| Cyflumetofen | 11496052 | 1 | [141095910] | [] | [141095910] | [] | 0 | Acaricides |
| Tau-Fluvalinate | 91768 | 2 | [2316093, 2316094] | [] | [2316093, 2316094] | [] | 0 | Pyrethroid |
| Azinphos-Ethyl | 17531 | 0 | [] | [] | [] | [] | 0 | Organophosphorus |
| Isoxathion | 29307 | 10 | [70318, 154897, 13608626, 13608628, 208501, 13608629, 13608631, 13608632, 13608633, 40572478] | [] | [70318, 154897, 13608626, 13608628, 208501, 13608629, 13608631, 13608632, 13608633, 40572478] | [] | 0 | Organophosphorus |
| Chlorfenapyr | 91778 | 7 | [18779853, 15239088, 15010164, 54456436, 11724470, 20093620, 67839709] | [] | [18779853, 15239088, 15010164, 54456436, 11724470, 20093620, 67839709] | [] | 0 | Others |
| Acequinocyl | 93315 | 12 | [93315, 4343367, 14672776, 21330062, 21330063, 102024497, 21330067, 4063412, 71333527, 21325241, 21325243, 20360830] | [93315, 14672776] | [4343367, 21330062, 21330063, 102024497, 21330067, 4063412, 71333527, 21325241, 21325243, 20360830] | ['93315>>139595618', '93315>>14672776'] | 2 | Acaricides |
| Fenobucarb | 19588 | 22 | [43522, 53437069, 42642, 12318874, 12318876, 12318878, 12318889, 21548843, 12318893, 12318895, 12318896, 14670271, 6984, 154220361, 14196819, 129651815, 131855976, 53859562, 17517, 92015598, 92015599, 71399674] | [] | [43522, 53437069, 42642, 12318874, 12318876, 12318878, 12318889, 21548843, 12318893, 12318895, 12318896, 14670271, 6984, 154220361, 14196819, 129651815, 131855976, 53859562, 17517, 92015598, 92015599, 71399674] | [] | 0 | Carbamate |
| Dicarbasulf | 6386307 | 1 | [6510008] | [] | [6510008] | [] | 0 | Carbamate |
| Thiocyclam | 35970 | 2 | [87894141, 88367630] | [] | [87894141, 88367630] | [] | 0 | Nereistoxin |
| 2-(Octylthio)ethanol | 19079 | 0 | [] | [] | [] | [] | 0 | Others |
| Bensultap | 87176 | 1 | [15710262] | [] | [15710262] | [] | 0 | Nereistoxin |
| Deltamethrin | 40585 | 10 | [89549635, 89179589, 101628462, 146674224, 146674225, 146674226, 6455384, 6455385, 6455386, 101627865] | [6455385] | [89549635, 89179589, 101628462, 146674224, 146674225, 146674226, 6455384, 6455386, 101627865] | ['40585>>6455385'] | 1 | Pyrethroid |
| Furathiocarb | 47759 | 8 | [47756, 47757, 47759, 18954032, 47761, 47762, 47763, 47760] | [] | [47756, 47757, 47759, 18954032, 47761, 47762, 47763, 47760] | [] | 0 | Carbamate |
| Xylylcarb | 17040 | 19 | [76334722, 3027978, 522251, 14915470, 14915471, 14915472, 42644, 19873, 134113700, 185656, 17592, 130011082, 7249, 186072, 12636, 130011105, 130011106, 23273850, 154109694] | [7249] | [76334722, 3027978, 522251, 14915470, 14915471, 14915472, 42644, 19873, 134113700, 185656, 17592, 130011082, 186072, 12636, 130011105, 130011106, 23273850, 154109694] | [] | 0 | Carbamate |
| Fenothiocarb | 44178 | 35 | [21993218, 13276419, 13276420, 13276422, 520713, 13276427, 1550859, 13276430, 13276431, 19845393, 13276435, 294549, 13228569, 13228570, 36691098, 13276447, 11392287, 71333032, 71333033, 16298, 13313449, 127262259, 74558, 20318920, 58892622, 517985, 9859302, 14311, 68132335, 39245295, 4438129, 71334385, 58963962, 59771, 3293437] | [] | [21993218, 13276419, 13276420, 13276422, 520713, 13276427, 1550859, 13276430, 13276431, 19845393, 13276435, 294549, 13228569, 13228570, 36691098, 13276447, 11392287, 71333032, 71333033, 16298, 13313449, 127262259, 74558, 20318920, 58892622, 517985, 9859302, 14311, 68132335, 39245295, 4438129, 71334385, 58963962, 59771, 3293437] | [] | 0 | Acaricides |
| (E)-1,3-Dichloropropene | 24726 | 4 | [93058, 287955, 560790, 643775] | [93058, 287955, 560790, 643775] | [] | ['24726>>287955'] | 1 | Nematicides |
| Endosulfan | 3224 | 5 | [74109123, 6431685, 13940, 120890, 54027199] | [13940, 120890] | [74109123, 6431685, 54027199] | ['3224>>120890', '3224>>13940'] | 2 | Organochlorine |
| Alanycarb | 9576091 | 3 | [21923138, 21923139, 88704971] | [] | [21923138, 21923139, 88704971] | [] | 0 | Carbamate |
| 3,5-Xylyl Methylcarbamate | 17563 | 14 | [76309313, 28617, 3027979, 7948, 12653, 25550, 17039, 14915473, 14915474, 185650, 688148, 42645, 6452059, 102239423] | [7948] | [76309313, 28617, 3027979, 12653, 25550, 17039, 14915473, 14915474, 185650, 688148, 42645, 6452059, 102239423] | [] | 0 | Carbamate |
| Imidacloprid | 86287518 | 23 | [135438466, 136244997, 137177223, 165363601, 101618973, 20114087, 9815679, 15390532, 13920968, 136264398, 136240208, 136397273, 136224125, 10130527, 135833314, 14673000, 136020079, 136866805, 183033, 136456570, 136456571, 136241533, 122226815] | [137177223, 20114087, 9815679, 15390532, 10130527, 136866805, 183033, 136241533, 122226815] | [135438466, 136244997, 165363601, 101618973, 13920968, 136264398, 136240208, 136397273, 136224125, 135833314, 14673000, 136020079, 136456570, 136456571] | ['86287518>>136241533', '86287518>>137177223', '86287518>>183033'] | 3 | Neonicotinoid |
| Phenothrin | 4767 | 6 | [13092066, 66896837, 78173382, 66896839, 66896650, 88777500] | [] | [13092066, 66896837, 78173382, 66896839, 66896650, 88777500] | [] | 0 | Pyrethroid |
| Esfenvalerate | 10342051 | 6 | [12922480, 92855793, 12922481, 56618163, 15390455, 9837437] | [] | [12922480, 92855793, 12922481, 56618163, 15390455, 9837437] | [] | 0 | Pyrethroid |
| Dicrotophos | 5371560 | 5 | [6444322, 5371562, 6443056, 21118675, 88713433] | [] | [6444322, 5371562, 6443056, 21118675, 88713433] | [] | 0 | Organophosphorus |
| Monocrotophos | 5371562 | 6 | [6444512, 6365306, 22842599, 5371560, 6443054, 151869754] | [] | [6444512, 6365306, 22842599, 5371560, 6443054, 151869754] | [] | 0 | Organophosphorus |
| Chlorpyrifos | 2730 | 6 | [181829, 85945287, 21804, 85838766, 13363034, 21249917] | [21804] | [181829, 85945287, 85838766, 13363034, 21249917] | ['2730>>21804'] | 1 | Organophosphorus |
| Nitenpyram | 3034287 | 7 | [9882947, 22450412, 6525197, 15132110, 3034287, 22450518, 87082615] | [] | [9882947, 22450412, 6525197, 15132110, 3034287, 22450518, 87082615] | [] | 0 | Neonicotinoid |
| Allethrins | 11442 | 8 | [14345093, 14345097, 11083, 141032749, 14345073, 14345075, 14345081, 14345087] | [] | [14345093, 14345097, 11083, 141032749, 14345073, 14345075, 14345081, 14345087] | [] | 0 | Pyrethroid |
| Phorate | 4790 | 5 | [21612128, 153783559, 17424, 17425, 9274] | [17424, 17425, 9274] | [21612128, 153783559] | ['4790>>17424', '4790>>9274'] | 2 | Organophosphorus |
| Empenthrin | 6434488 | 3 | [13787817, 12907802, 13265795] | [] | [13787817, 12907802, 13265795] | [] | 0 | Pyrethroid |
| Phosalone | 4793 | 1 | [207757] | [] | [207757] | [] | 0 | Organophosphorus |
| Pyridaphenthion | 8381 | 10 | [25764994, 13180002, 13180003, 13180005, 13180006, 13180007, 21418089, 13180015, 6454521, 74333] | [] | [25764994, 13180002, 13180003, 13180005, 13180006, 13180007, 21418089, 13180015, 6454521, 74333] | [] | 0 | Organophosphorus |
| [(E)-3-methylsulfonylbutan-2-ylideneamino] N-methylcarbamate | 9571009 | 1 | [20195606] | [] | [20195606] | [] | 0 | Carbamate |
| Spirodiclofen | 177863 | 3 | [54719923, 142670565, 11304039] | [] | [54719923, 142670565, 11304039] | [] | 0 | Acaricides |
| Indoxacarb | 107720 | 7 | [69540130, 139594186, 139597707, 149549437, 69344719, 155884400, 125340765] | [139594186, 139597707, 155884400] | [69540130, 149549437, 69344719, 125340765] | ['107720>>139594186', '107720>>139597707', '107720>>155884400'] | 3 | Others |
| (Z)-Metaflumizone | 9827529 | 1 | [89251036] | [] | [89251036] | [] | 0 | Others |
| Prallethrin(Mixture of diastereomers) | 9839306 | 2 | [11442, 12918206] | [] | [11442, 12918206] | [] | 0 | Pyrethroid |
| Spiromesifen | 9907412 | 2 | [16752776, 54719924] | [] | [16752776, 54719924] | [] | 0 | Others |
| Oxythioquinox | 17109 | 1 | [20468957] | [] | [20468957] | [] | 0 | Acaricides |
| Ethion | 3286 | 0 | [] | [] | [] | [] | 0 | Organophosphorus |
| Hexachlorocyclohexane | 727 | 3 | [441093, 15691, 12914037] | [441093, 15691, 12914037] | [] | ['727>>1038\|12914037\|312', '727>>1038\|312\|441093', '727>>15691', '727>>441093'] | 4 | Organochlorine |
| demeton-S-methyl | 13526 | 6 | [116226, 4618, 54102541, 189264, 24723, 28213] | [] | [116226, 4618, 54102541, 189264, 24723, 28213] | [] | 0 | Organophosphorus |
| Ethoprophos | 3289 | 7 | [22119594, 11831182, 182032, 13302641, 19096563, 156276, 13362995] | [182032] | [22119594, 11831182, 13302641, 19096563, 156276, 13362995] | ['3289>>182032'] | 1 | Organophosphorus |
| Azinphos-methyl | 2268 | 7 | [71412256, 71414081, 57169476, 13746, 135372946, 17531, 71412255] | [] | [71412256, 71414081, 57169476, 13746, 135372946, 17531, 71412255] | [] | 0 | Organophosphorus |
| Trichlorfon | 5853 | 16 | [12673281, 20454, 156008968, 85966216, 164746, 3013932, 13134, 31343, 92144, 114513, 13717714, 99092, 14577078, 12550616, 22235, 5853] | [3013932, 13134, 114513, 5853] | [12673281, 20454, 156008968, 85966216, 164746, 31343, 92144, 13717714, 99092, 14577078, 12550616, 22235] | ['5853>>114513', '5853>>114636', '5853>>13134', '5853>>176943', '5853>>3013932', '5853>>3039'] | 6 | Organophosphorus |
| Pirimiphos-methyl | 34526 | 7 | [13369988, 165362405, 9966798, 190609, 135448147, 31957, 13789821] | [] | [13369988, 165362405, 9966798, 190609, 135448147, 31957, 13789821] | [] | 0 | Organophosphorus |
| Fenpropathrin | 47326 | 3 | [13386368, 146881726, 90808078] | [] | [13386368, 146881726, 90808078] | [] | 0 | Pyrethroid |
| Famphur | 5859 | 10 | [3048774, 200999, 120174, 54190608, 200670, 84760, 668793, 201149, 201150, 70399] | [] | [3048774, 200999, 120174, 54190608, 200670, 84760, 668793, 201149, 201150, 70399] | [] | 0 | Organophosphorus |
| Transfluthrin | 656612 | 7 | [54322177, 53878468, 90392325, 2734029, 140309334, 140309340, 140309342] | [] | [54322177, 53878468, 90392325, 2734029, 140309334, 140309340, 140309342] | [] | 0 | Pyrethroid |
| Bifenazate | 176879 | 10 | [11312193, 171042086, 139595846, 171042088, 139596138, 22020266, 69250380, 123550961, 85699030, 139594559] | [139595846, 139596138, 69250380, 139594559] | [11312193, 171042086, 171042088, 22020266, 123550961, 85699030] | ['176879>>139594559', '176879>>139595846', '176879>>139596138', '176879>>69250380'] | 4 | Acaricides |
| Diethoxy-[ethoxy(methoxymethyl)phosphinothioyl]oxy-thioxo-$l^{5}-phosphane | 53297399 | 0 | [] | [] | [] | [] | 0 | Organophosphorus |
| Thiometon | 12541 | 10 | [88482, 525385, 57254826, 59755053, 3118, 75917, 53958772, 13526, 101416438, 59154905] | [3118] | [88482, 525385, 57254826, 59755053, 75917, 53958772, 13526, 101416438, 59154905] | [] | 0 | Organophosphorus |
| Acrinathrin | 6436606 | 4 | [149435968, 161829869, 87657941, 13093798] | [] | [149435968, 161829869, 87657941, 13093798] | [] | 0 | Pyrethroid |
| alpha-Chlorfenvinphos | 5377791 | 6 | [13376386, 21117003, 24841683, 13084215, 138402105, 6445756] | [] | [13376386, 21117003, 24841683, 13084215, 138402105, 6445756] | [] | 0 | Organophosphorus |
| Spinetoram | 53297414 | 0 | [] | [] | [] | [] | 0 | Others |
| Bendiocarb | 2314 | 6 | [31522, 21400649, 62765, 21400657, 31517, 145738782] | [62765] | [31522, 21400649, 21400657, 31517, 145738782] | ['2314>>62765'] | 1 | Carbamate |
| Diafenthiuron | 3034380 | 5 | [101135180, 138388115, 13272150, 13272155, 13272156] | [] | [101135180, 138388115, 13272150, 13272155, 13272156] | [] | 0 | Acaricides |
| Silafluofen | 92430 | 0 | [] | [] | [] | [] | 0 | Pyrethroid |
| Temephos | 5392 | 4 | [580328, 70897, 132275285, 101694728] | [] | [580328, 70897, 132275285, 101694728] | [] | 0 | Organophosphorus |
| Fenthion | 3346 | 9 | [23046, 185099, 597806, 66128, 18391, 74390, 23207414, 19577, 19578] | [23046, 19577, 19578] | [185099, 597806, 66128, 18391, 74390, 23207414] | ['3346>>23046'] | 1 | Organophosphorus |
| Fenvalerate | 3347 | 14 | [139652548, 188070, 157223, 12922479, 20275888, 182481, 12895088, 13090449, 188116, 100965877, 184823, 181912, 143074969, 13326427] | [188070, 188116, 181912] | [139652548, 157223, 12922479, 20275888, 182481, 12895088, 13090449, 100965877, 184823, 143074969, 13326427] | ['3347>>188070', '3347>>188116'] | 2 | Pyrethroid |
| Pyraclofos | 93460 | 5 | [12901121, 12901092, 12901128, 101647787, 12901119] | [] | [12901121, 12901092, 12901128, 101647787, 12901119] | [] | 0 | Organophosphorus |
| (1E)-2-methyl-2-(methylthio)propanal O-[(methylamino)carbonyl]oxime | 9570071 | 5 | [9576932, 133625927, 9570092, 9570093, 9568700] | [] | [9576932, 133625927, 9570092, 9570093, 9568700] | [] | 0 | Carbamate |
| Fipronil | 3352 | 17 | [22618497, 23079074, 23079075, 22673275, 137353829, 11784549, 22618634, 165363563, 11430864, 15456945, 86613234, 9953940, 155907320, 51066521, 9933690, 3078139, 22618429] | [22618497, 23079074, 23079075, 22673275, 15456945, 86613234, 9953940, 9933690, 3078139] | [137353829, 11784549, 22618634, 165363563, 11430864, 155907320, 51066521, 22618429] | ['3352>>15456945', '3352>>22618497', '3352>>22673275', '3352>>23079074', '3352>>23079075', '3352>>3078139', '3352>>86613234', '3352>>9933690', '3352>>9953940', '9953940>>3352'] | 10 | Pyrazole |
| Methomyl | 5353758 | 9 | [133681536, 5491811, 5365286, 5491367, 5492999, 9570633, 19884468, 6165177, 55255295] | [] | [133681536, 5491811, 5365286, 5491367, 5492999, 9570633, 19884468, 6165177, 55255295] | [] | 0 | Carbamate |
| Triazamate | 86306 | 2 | [14299898, 21988797] | [14299898] | [21988797] | ['86306>>14299898'] | 1 | Others |
| Flucythrinate | 50980 | 7 | [20487624, 16722122, 13090480, 57241649, 57293299, 57179676, 57127069] | [] | [20487624, 16722122, 13090480, 57241649, 57293299, 57179676, 57127069] | [] | 0 | Pyrethroid |
| Chlorpyrifos-methyl | 21803 | 9 | [85859587, 85859588, 23017, 2730, 17915276, 21805, 92279, 23220151, 150816666] | [23017, 2730] | [85859587, 85859588, 17915276, 21805, 92279, 23220151, 150816666] | ['21803>>2730'] | 1 | Organophosphorus |
| Hostaquick | 62773 | 1 | [23274165] | [] | [23274165] | [] | 0 | Organophosphorus |
| Coumaphos | 2871 | 4 | [154517704, 10496498, 9453, 5355079] | [] | [154517704, 10496498, 9453, 5355079] | [] | 0 | Acaricides |
| Azamethiphos | 71482 | 0 | [] | [] | [] | [] | 0 | Organophosphorus |
| [(E)-3-methylsulfanylbutan-2-ylideneamino] N-methylcarbamate | 5360962 | 4 | [5745504, 9571009, 133681419, 9576739] | [] | [5745504, 9571009, 133681419, 9576739] | [] | 0 | Carbamate |
| Methacrifos | 3034435 | 3 | [153915378, 87566053, 3035167] | [] | [153915378, 87566053, 3035167] | [] | 0 | Organophosphorus |
| Naled | 4420 | 5 | [154253558, 154253559, 21398776, 21314683, 3039] | [3039] | [154253558, 154253559, 21398776, 21314683] | ['4420>>3039'] | 1 | Organophosphorus |
| Propargite | 4936 | 3 | [16033, 15801298, 14678926] | [] | [16033, 15801298, 14678926] | [] | 0 | Acaricides |
| Tebupirimfos | 93516 | 4 | [149251600, 22243747, 163499, 86278326] | [] | [149251600, 22243747, 163499, 86278326] | [] | 0 | Organophosphorus |
| Propoxur | 4944 | 13 | [13277697, 77539, 197892, 23274309, 24840806, 82537, 91431723, 38097, 280786, 25777, 20949, 21279414, 140422422] | [] | [13277697, 77539, 197892, 23274309, 24840806, 82537, 91431723, 38097, 280786, 25777, 20949, 21279414, 140422422] | [] | 0 | Carbamate |
| Tebufenpyrad | 86354 | 8 | [25137220, 7010409, 14460816, 59678674, 14460821, 139598008, 59678617, 58503101] | [139598008, 59678617] | [25137220, 7010409, 14460816, 59678674, 14460821, 58503101] | ['86354>>139598008', '86354>>59678617'] | 2 | Pyrazole |
| Fenazaquin | 86356 | 17 | [86178016, 45691520, 139594723, 59678659, 86178021, 86178023, 141905642, 86178027, 139597870, 59678639, 23196560, 67620305, 23196528, 67620212, 67619861, 59678619, 86178015] | [45691520, 139594723, 139597870, 59678639] | [86178016, 59678659, 86178021, 86178023, 141905642, 86178027, 23196560, 67620305, 23196528, 67620212, 67619861, 59678619, 86178015] | ['86356>>139594723', '86356>>139597870', '86356>>45691520', '86356>>59678639'] | 4 | Acaricides |
| Mevinphos | 5355863 | 2 | [6381336, 22821767] | [] | [6381336, 22821767] | [] | 0 | Organophosphorus |
| (1S)-cis-(alphaR)-cypermethrin | 3086172 | 4 | [156232, 169436650, 100899213, 117956754] | [] | [156232, 169436650, 100899213, 117956754] | [] | 0 | Pyrethroid |
| Fenamiphos | 31070 | 4 | [36028, 36027, 68762852, 152677686] | [] | [36028, 36027, 68762852, 152677686] | [] | 0 | Phosphoramido |
| Cypermethrin | 2912 | 17 | [169451428, 15687973, 182438, 12795783, 78173324, 78173325, 12810924, 14330639, 21829872, 12873617, 102063091, 21829877, 12822232, 143550266, 5311131, 56688284, 75951486] | [182438] | [169451428, 15687973, 12795783, 78173324, 78173325, 12810924, 14330639, 21829872, 12873617, 102063091, 21829877, 12822232, 143550266, 5311131, 56688284, 75951486] | ['2912>>182438'] | 1 | Pyrethroid |
| CID 9809914 | 9809914 | 1 | [12874360] | [] | [12874360] | [] | 0 | Pyrethroid |
| Chlordane, technical | 5993 | 2 | [14029296, 19519] | [] | [14029296, 19519] | [] | 0 | Organochlorine |
| Etoxazole | 153974 | 4 | [138394724, 24848458, 149787507, 148367204] | [138394724, 24848458] | [149787507, 148367204] | ['153974>>138394724', '153974>>24848458'] | 2 | Acaricides |
| Methiocarb | 16248 | 7 | [154332131, 21279429, 16589, 12889680, 17521, 81853, 138394719] | [16589, 17521, 81853] | [154332131, 21279429, 12889680, 138394719] | ['16248>>16589', '16248>>17521', '16248>>81853'] | 3 | Carbamate |
| Formetanate | 31099 | 3 | [117570, 21279486, 44152782] | [] | [117570, 21279486, 44152782] | [] | 0 | Acaricides |
| Profenofos | 38779 | 9 | [102156261, 190054, 101749926, 13368229, 85800838, 86016019, 15400052, 85800889, 101213978] | [] | [102156261, 190054, 101749926, 13368229, 85800838, 86016019, 15400052, 85800889, 101213978] | [] | 0 | Organophosphorus |
| Omethoate | 14210 | 5 | [88842856, 21718857, 85741422, 169561, 213467] | [] | [88842856, 21718857, 85741422, 169561, 213467] | [] | 0 | Organophosphorus |
| Permethrin | 40326 | 8 | [139840870, 156028136, 13034991, 12799248, 152567638, 26295, 53959928, 190015] | [26295] | [139840870, 156028136, 13034991, 12799248, 152567638, 53959928, 190015] | [] | 0 | Pyrethroid |
| Fluacrypyrim | 9954185 | 4 | [70233376, 68515610, 163418352, 11223662] | [] | [70233376, 68515610, 163418352, 11223662] | [] | 0 | Acaricides |
| Cyphenothrin | 38283 | 6 | [13092064, 93218, 12822244, 67629412, 70626646, 78173142] | [93218] | [13092064, 12822244, 67629412, 70626646, 78173142] | [] | 0 | Pyrethroid |
| Methidathion | 13709 | 6 | [3084322, 88197, 57146823, 53723883, 190999, 54024921] | [] | [3084322, 88197, 57146823, 53723883, 190999, 54024921] | [] | 0 | Organophosphorus |
| 3,3-Dimethyl-1-(methylthio)-2-butanone O-(methylcarbamoyl)oxime | 5361043 | 7 | [21565507, 21565509, 21565511, 5491759, 5491760, 5361040, 21318204] | [] | [21565507, 21565509, 21565511, 5491759, 5491760, 5361040, 21318204] | [] | 0 | Carbamate |
| Fenoxycarb | 51605 | 19 | [13520782, 54553614, 14098199, 129829018, 18966691, 22119590, 85803431, 19881385, 19881386, 54348478, 13374527, 18966594, 13254, 129859418, 85803483, 15766499, 11031651, 19424114, 4676725] | [129829018, 11031651, 4676725] | [13520782, 54553614, 14098199, 18966691, 22119590, 85803431, 19881385, 19881386, 54348478, 13374527, 18966594, 13254, 129859418, 85803483, 15766499, 19424114] | ['51605>>11031651', '51605>>129829018', '51605>>4676725'] | 3 | Carbamate |
| Ethanimidothioic acid, 2-(dimethylamino)-N-(((methylamino)carbonyl)oxy)-2-oxo-, methyl ester | 9595287 | 7 | [23460547, 13131877, 13093843, 88483957, 6399255, 13093849, 24837947] | [] | [23460547, 13131877, 13093843, 88483957, 6399255, 13093849, 24837947] | [] | 0 | Carbamate |
| Pirimicarb | 31645 | 9 | [146037633, 185602, 135420611, 129736354, 15390533, 67885285, 100945355, 93139, 182004] | [146037633, 185602, 135420611, 93139, 182004] | [129736354, 15390533, 67885285, 100945355] | ['31645>>135420611', '31645>>146037633', '31645>>182004', '31645>>185602', '31645>>93139'] | 5 | Carbamate |
| Methyl isothiocyanate | 11167 | 2 | [19035076, 20692359] | [] | [19035076, 20692359] | [] | 0 | Nematicides |
| Flubendiamide | 11193251 | 3 | [165362497, 101853051, 169444507] | [] | [165362497, 101853051, 169444507] | [] | 0 | Others |
| Malathion | 4004 | 12 | [14461667, 225316, 85581062, 90952615, 20337862, 86097653, 50904182, 15415, 580760, 6913625, 57579354, 3032927] | [20337862, 15415, 6913625] | [14461667, 225316, 85581062, 90952615, 86097653, 50904182, 580760, 57579354, 3032927] | ['4004>>15415', '4004>>20337862', '4004>>6913625'] | 3 | Organophosphorus |
| Spirotetramat | 9969573 | 12 | [139596673, 54708610, 139594401, 138394788, 71312325, 25135301, 10179592, 139597999, 139594160, 150675826, 59229527, 68416382] | [139596673, 54708610, 139594401, 138394788, 71312325, 139597999, 139594160] | [25135301, 10179592, 150675826, 59229527, 68416382] | ['9969573>>138394788', '9969573>>139594160', '9969573>>139594401', '9969573>>139596673', '9969573>>139597999', '9969573>>54708610', '9969573>>71312325'] | 7 | Others |
| Carbosulfan | 41384 | 4 | [41384, 154700115, 18347195, 187367] | [41384, 154700115] | [18347195, 187367] | ['41384>>154700115'] | 1 | Carbamate |
| Ethiprole | 9930667 | 3 | [22618433, 10949642, 86278308] | [22618433] | [10949642, 86278308] | ['9930667>>22618433'] | 1 | Pyrazole |
| Triazophos | 32184 | 5 | [85996835, 85996836, 40572490, 77910, 69933851] | [77910] | [85996835, 85996836, 40572490, 69933851] | [] | 0 | Organophosphorus |
| Resmethrin | 5053 | 2 | [12364396, 14258462] | [] | [12364396, 14258462] | [] | 0 | Pyrethroid |
| Acephate | 1982 | 3 | [4096, 85866905, 85866904] | [4096] | [85866905, 85866904] | ['1982>>4096'] | 1 | Phosphoramido |
| Pyrimidifen | 6451139 | 2 | [141905718, 18354214] | [] | [141905718, 18354214] | [] | 0 | Acaricides |
| Diazinon | 3017 | 8 | [91412032, 129682113, 186178, 190308, 54549926, 592775, 135444498, 13754] | [13754] | [91412032, 129682113, 186178, 190308, 54549926, 592775, 135444498] | ['3017>>13754'] | 1 | Organophosphorus |
| Ethiofencarb | 34766 | 6 | [58048, 119490, 3035207, 13434024, 140995819, 522167] | [] | [58048, 119490, 3035207, 13434024, 140995819, 522167] | [] | 0 | Carbamate |
| Thiamethoxam | 5821911 | 0 | [] | [] | [] | [] | 0 | Neonicotinoid |
| Chlorantraniliprole | 11271640 | 16 | [135564546, 68396004, 167138408, 68396009, 10115786, 164597901, 129078830, 154578288, 11317968, 154366004, 155666902, 169433942, 165361848, 90367673, 165361850, 169435964] | [135564546, 10115786, 90367673] | [68396004, 167138408, 68396009, 164597901, 129078830, 154578288, 11317968, 154366004, 155666902, 169433942, 165361848, 165361850, 169435964] | ['11271640>>10115786', '11271640>>135564546', '11271640>>90367673'] | 3 | Pyrazole |
| Hexythiazox | 13218777 | 2 | [70513419, 141229372] | [] | [70513419, 141229372] | [] | 0 | Acaricides |
| Clofenotane | 3036 | 7 | [3035, 101333683, 85774004, 6294, 101307, 441117, 441118] | [3035, 6294, 441117, 441118] | [101333683, 85774004, 101307] | ['3036>>1038\|3035\|312', '3036>>3035', '3036>>441117', '3036>>6294'] | 4 | Organochlorine |
| Fenpyroximate | 9576412 | 10 | [21988800, 14300736, 14300737, 165363497, 14300682, 14300724, 14300731, 14300732, 14300734, 14300735] | [] | [21988800, 14300736, 14300737, 165363497, 14300682, 14300724, 14300731, 14300732, 14300734, 14300735] | [] | 0 | Acaricides |
| Cyfluthrin | 104926 | 8 | [74326725, 13150406, 13150410, 13150411, 102063092, 10377109, 13092340, 13335384] | [] | [74326725, 13150406, 13150410, 13150411, 102063092, 10377109, 13092340, 13335384] | [] | 0 | Pyrethroid |
| Dichlorvos | 3039 | 7 | [66145, 86500, 6597, 114636, 85613422, 176943, 175423960] | [6597, 114636, 176943] | [66145, 86500, 85613422, 175423960] | ['3039>>114636', '3039>>6597'] | 2 | Organophosphorus |
| Pentachlorophenol | 992 | 29 | [6914, 6028, 131876757, 21013, 15767, 66603, 13618, 11571, 13619, 78177980, 88095421, 27582, 54302400, 13636, 11334, 51373383, 14537, 11855, 11859, 119254, 53703129, 21119838, 992, 85816035, 152804, 92261, 6899, 220151, 7933] | [6914, 66603, 13618, 13619, 27582, 11334, 14537, 11855, 11859, 992, 6899, 7933] | [6028, 131876757, 21013, 15767, 11571, 78177980, 88095421, 54302400, 13636, 51373383, 119254, 53703129, 21119838, 85816035, 152804, 92261, 220151] | ['11855>>992', '8370>>992', '992>>66603'] | 3 | Organochlorine |
| Fenitrothion | 31200 | 19 | [17412, 3053970, 123163, 15663943, 167496, 127435, 18788052, 3039448, 85822809, 3039450, 165212, 31200, 79330, 16738, 41194, 198380, 25843, 4414327, 56841722] | [3053970, 165212, 31200, 79330, 16738, 25843] | [17412, 123163, 15663943, 167496, 127435, 18788052, 3039448, 85822809, 3039450, 41194, 198380, 4414327, 56841722] | ['31200>>139596254', '31200>>165212', '31200>>16738', '31200>>25843', '31200>>3053970', '31200>>79330'] | 6 | Organophosphorus |
| Parathion | 991 | 13 | [85843561, 85843562, 36106, 201452, 13041322, 21447693, 132561421, 9395, 991, 220, 15836381, 86176062, 200990] | [9395, 991, 220] | [85843561, 85843562, 36106, 201452, 13041322, 21447693, 132561421, 15836381, 86176062, 200990] | ['962\|991>>1038\|3683036\|644235', '991>>220', '991>>9395'] | 3 | Organophosphorus |
| Chlormephos | 32739 | 0 | [] | [] | [] | [] | 0 | Organophosphorus |
| Amitraz | 36324 | 2 | [20187603, 143167325] | [] | [20187603, 143167325] | [] | 0 | Acaricides |
| Sulfluramid | 77797 | 8 | [10507011, 111914, 14496522, 74483, 88321907, 101895126, 69785, 91030301] | [10507011, 14496522, 74483, 101895126, 69785] | [111914, 88321907, 91030301] | ['77797>>101895126', '77797>>10507011', '77797>>14496522', '77797>>69785', '77797>>74483'] | 5 | Others |
| Carbaryl | 6129 | 31 | [14344, 22752778, 151418515, 135956, 259741, 54505377, 56842660, 177830, 56842676, 153894325, 20090231, 21956, 6451525, 23493, 17481, 20090321, 68451284, 23618388, 7005, 21341, 185565, 21360480, 21342, 15148256, 24840810, 79338, 36594, 101699447, 101699448, 101699449, 165355900] | [21956, 7005, 21341, 21342] | [14344, 22752778, 151418515, 135956, 259741, 54505377, 56842660, 177830, 56842676, 153894325, 20090231, 6451525, 23493, 17481, 20090321, 68451284, 23618388, 185565, 21360480, 15148256, 24840810, 79338, 36594, 101699447, 101699448, 101699449, 165355900] | ['1038\|6129\|962>>280\|644041\|7005', '6129>>21341', '6129>>21342', '6129>>21956', '6129>>7005'] | 5 | Carbamate |
| Phoxim | 9570290 | 2 | [9570291, 9562484] | [] | [9570291, 9562484] | [] | 0 | Organophosphorus |
| Metolcarb | 14322 | 25 | [14915469, 17040, 154346640, 42646, 17563, 36380, 101073310, 154204027, 21409338, 130011067, 130011068, 129736381, 134097724, 23161405, 6452033, 21680069, 154101574, 154227792, 342, 23273692, 235375, 167794, 131858933, 75765, 130011259] | [342] | [14915469, 17040, 154346640, 42646, 17563, 36380, 101073310, 154204027, 21409338, 130011067, 130011068, 129736381, 134097724, 23161405, 6452033, 21680069, 154101574, 154227792, 23273692, 235375, 167794, 131858933, 75765, 130011259] | [] | 0 | Carbamate |
| Azocyclotin | 91634 | 0 | [] | [] | [] | [] | 0 | Acaricides |
| Propetamphos | 5372405 | 2 | [88478628, 101318351] | [] | [88478628, 101318351] | [] | 0 | Phosphoramido |
| Milbemycin A3 | 9828343 | 8 | [76334273, 76319811, 76871849, 10415018, 24748203, 6440785, 9828343, 10554552] | [] | [76334273, 76319811, 76871849, 10415018, 24748203, 6440785, 9828343, 10554552] | [] | 0 | Others |
| Nicotine | 89594 | 60 | [211457, 854019, 11991300, 28064517, 11401098, 68107, 14286224, 181139, 62228, 10261653, 59590422, 148819607, 409, 13945370, 25203099, 412, 413, 415, 53919010, 115108, 431, 432, 435, 12749620, 437, 6453941, 439, 58537397, 436, 145285561, 11513911, 10613308, 3015101, 23615292, 438, 14272, 26248770, 91462, 88903, 3035848, 16082121, 98464332, 68602291, 25201488, 928594, 12000213, 12159574, 12960087, 439383, 12960088, 74074, 102579424, 71751013, 11564518, 71414247, 10633832, 108, 114, 10352249, 10219774] | [854019, 68107, 181139, 62228, 148819607, 409, 25203099, 412, 413, 415, 53919010, 115108, 431, 432, 435, 437, 439, 436, 3015101, 23615292, 438, 14272, 91462, 3035848, 25201488, 439383, 108, 114, 10219774] | [211457, 11991300, 28064517, 11401098, 14286224, 10261653, 59590422, 13945370, 12749620, 6453941, 58537397, 145285561, 11513911, 10613308, 26248770, 88903, 16082121, 98464332, 68602291, 928594, 12000213, 12159574, 12960087, 12960088, 74074, 102579424, 71751013, 11564518, 71414247, 10633832, 10352249] | ['1038\|22833512\|89594\|977>>409\|5886\|962', '17473\|89594>>3035848\|6031', '89594>>1038\|431', '89594>>108', '89594>>114', '89594>>115108', '89594>>14272', '89594>>148819607', '89594>>181139', '89594>>23615292', '89594>>3015101', '89594>>3035848', '89594>>409', '89594>>412', '89594>>413', '89594>>415', '89594>>431', '89594>>432', '89594>>435', '89594>>436', '89594>>437', '89594>>438', '89594>>439', '89594>>53919010', '89594>>62228', '89594>>91462'] | 26 | Others |
| 2-[3-[2,6-Dichloro-4-(3,3-dichloroprop-2-enoxy)phenoxy]propoxy]-6-(trifluoromethyl)pyridine | 24847867 | 0 | [] | [] | [] | [] | 0 | Others |
| Fenbutatin oxide | 16683004 | 0 | [] | [] | [] | [] | 0 | Acaricides |

**SI Table 4:** **Descriptive Statistics of Transformation Products ADME_Tox Properties** shows the total number of unique transformation products generated (count; 21284), average value (mean), standard deviation (std), range of values (min, max), and three quartiles for each property. It lists key physicochemical properties like molecular weight, logarithm of the partition coefficient (logP), number of H-bond acceptor and donor atoms (hydrogen_bond_acceptors / hydrogen_bond_donors), topological polar surface area (tpsa), number of stereocenters in the molecule (stereo_centers), and predicted hydration free energy (HydrationFreeEnergy_FreeSolv). The dataset includes a wide range of descriptors that cover different pharmacological and toxicological areas. Under Drug-Likeness and Bioavailability, it includes compliance with Lipinski’s Rule of Five, the Quantitative Estimate of Drug-likeness (QED), predicted oral bioavailability (Bioavailability_Ma), volume of distribution at steady state (VDss_Lombardo), plasma protein binding ratio (PPBR_AZ), and predictions for aqueous solubility (Solubility_AqSolDB). The Metabolism and Enzyme Interactions section features predictions for substrate or inhibitor activity across various cytochrome P450 enzymes, such as CYP1A2_Veith, CYP2D6_Veith, and CYP3A4_Substrate_CarbonMangels. In the Toxicity and Safety section, the dataset provides results from the Ames mutagenicity test (AMES), predictions for carcinogenicity (Carcinogens_Lagunin), clinical toxicity (ClinTox), and potential for drug-induced liver injury (DILI). It also includes indicators for skin sensitization and cardiac toxicity via hERG inhibition, as well as predicted lethal dose (LD50_Zhu). Absorption and Permeability metrics cover blood-brain barrier permeability (BBB_Martins), human intestinal absorption (HIA_Hou), and cell permeability predictions from Caco2_Wang and PAMPA_NCATS, along with predictions for P-glycoprotein substrates (Pgp_Broccatelli). The dataset also includes Nuclear Receptor and Stress Response Pathways, with predictions for nuclear receptor binding (e.g., NR-AR, NR-ER, NR-AhR, NR-PPAR-gamma) and activation of stress response pathways (e.g., SR-ARE, SR-HSE, SR-MMP, SR-p53). Finally, Clearance and Half-Life properties feature predictions for hepatic and microsomal clearance (Clearance_Hepatocyte_AZ, Clearance_Microsome_AZ) and the predicted half-life (Half_Life_Obach) of each compound.

| ADME Tox Property | count | mean | std | min | 25% | 50% | 75% | max |
| --- | --- | --- | --- | --- | --- | --- | --- | --- |
| molecular_weight | 19392 | 563.7 | 288.6 | 87.1 | 368.3 | 461.3 | 693.8 | 1665.8 |
| logP | 19392 | 6.6 | 5.4 | -3.7 | 3.1 | 5.0 | 7.9 | 32.1 |
| hydrogen_bond_acceptors | 19392 | 7.0 | 3.4 | 0.0 | 4.0 | 6.0 | 9.0 | 21.0 |
| hydrogen_bond_donors | 19392 | 1.1 | 1.1 | 0.0 | 0.0 | 1.0 | 2.0 | 7.0 |
| Lipinski | 19392 | 2.9 | 1.1 | 1.0 | 2.0 | 3.0 | 4.0 | 4.0 |
| QED | 19392 | 0.4 | 0.3 | 0.0 | 0.1 | 0.3 | 0.6 | 0.9 |
| stereo_centers | 19392 | 3.0 | 4.0 | 0.0 | 0.0 | 2.0 | 3.0 | 25.0 |
| tpsa | 19392 | 98.1 | 47.6 | 0.0 | 62.2 | 88.0 | 127.0 | 301.5 |
| AMES | 19392 | 0.2 | 0.2 | 0.0 | 0.1 | 0.2 | 0.4 | 1.0 |
| BBB_Martins | 19392 | 0.7 | 0.3 | 0.0 | 0.5 | 0.7 | 0.9 | 1.0 |
| Bioavailability_Ma | 19392 | 0.7 | 0.2 | 0.1 | 0.5 | 0.7 | 0.8 | 1.0 |
| CYP1A2_Veith | 19392 | 0.2 | 0.3 | 0.0 | 0.0 | 0.1 | 0.4 | 1.0 |
| CYP2C19_Veith | 19392 | 0.5 | 0.3 | 0.0 | 0.2 | 0.5 | 0.8 | 1.0 |
| CYP2C9_Substrate_CarbonMangels | 19392 | 0.2 | 0.2 | 0.0 | 0.0 | 0.2 | 0.3 | 0.8 |
| CYP2C9_Veith | 19392 | 0.4 | 0.3 | 0.0 | 0.1 | 0.3 | 0.7 | 1.0 |
| CYP2D6_Substrate_CarbonMangels | 19392 | 0.1 | 0.1 | 0.0 | 0.0 | 0.1 | 0.2 | 0.8 |
| CYP2D6_Veith | 19392 | 0.1 | 0.1 | 0.0 | 0.0 | 0.1 | 0.2 | 0.8 |
| CYP3A4_Substrate_CarbonMangels | 19392 | 0.7 | 0.2 | 0.0 | 0.6 | 0.8 | 0.8 | 1.0 |
| CYP3A4_Veith | 19392 | 0.5 | 0.3 | 0.0 | 0.2 | 0.5 | 0.7 | 1.0 |
| Carcinogens_Lagunin | 19392 | 0.3 | 0.2 | 0.0 | 0.1 | 0.3 | 0.4 | 1.0 |
| ClinTox | 19392 | 0.1 | 0.1 | 0.0 | 0.0 | 0.0 | 0.1 | 0.7 |
| DILI | 19392 | 0.6 | 0.3 | 0.0 | 0.3 | 0.6 | 0.8 | 1.0 |
| HIA_Hou | 19392 | 0.9 | 0.2 | 0.0 | 1.0 | 1.0 | 1.0 | 1.0 |
| NR-AR-LBD | 19392 | 0.0 | 0.0 | 0.0 | 0.0 | 0.0 | 0.0 | 0.3 |
| NR-AR | 19392 | 0.1 | 0.1 | 0.0 | 0.0 | 0.0 | 0.0 | 0.6 |
| NR-AhR | 19392 | 0.2 | 0.3 | 0.0 | 0.0 | 0.1 | 0.2 | 1.0 |
| NR-Aromatase | 19392 | 0.2 | 0.2 | 0.0 | 0.1 | 0.1 | 0.3 | 0.9 |
| NR-ER-LBD | 19392 | 0.1 | 0.1 | 0.0 | 0.0 | 0.0 | 0.1 | 0.9 |
| NR-ER | 19392 | 0.2 | 0.1 | 0.0 | 0.1 | 0.1 | 0.2 | 0.9 |
| NR-PPAR-gamma | 19392 | 0.1 | 0.1 | 0.0 | 0.0 | 0.0 | 0.1 | 0.8 |
| PAMPA_NCATS | 19392 | 0.8 | 0.2 | 0.0 | 0.7 | 0.9 | 1.0 | 1.0 |
| Pgp_Broccatelli | 19392 | 0.6 | 0.4 | 0.0 | 0.1 | 0.6 | 0.9 | 1.0 |
| SR-ARE | 19392 | 0.4 | 0.2 | 0.0 | 0.2 | 0.4 | 0.6 | 1.0 |
| SR-ATAD5 | 19392 | 0.0 | 0.0 | 0.0 | 0.0 | 0.0 | 0.0 | 0.8 |
| SR-HSE | 19392 | 0.1 | 0.1 | 0.0 | 0.0 | 0.1 | 0.1 | 0.9 |
| SR-MMP | 19392 | 0.4 | 0.3 | 0.0 | 0.1 | 0.4 | 0.7 | 1.0 |
| SR-p53 | 19392 | 0.1 | 0.1 | 0.0 | 0.0 | 0.0 | 0.1 | 0.9 |
| Skin_Reaction | 19392 | 0.5 | 0.2 | 0.0 | 0.3 | 0.5 | 0.6 | 1.0 |
| hERG | 19392 | 0.5 | 0.3 | 0.0 | 0.2 | 0.6 | 0.8 | 1.0 |
| Caco2_Wang | 19392 | -4.9 | 0.5 | -7.3 | -5.2 | -4.9 | -4.6 | -3.3 |
| Clearance_Hepatocyte_AZ | 19392 | 79.1 | 43.6 | -57.5 | 47.6 | 87.4 | 113.0 | 192.7 |
| Clearance_Microsome_AZ | 19392 | 72.5 | 42.2 | -42.7 | 39.9 | 74.1 | 105.2 | 183.4 |
| Half_Life_Obach | 19392 | 25.1 | 42.2 | -69.7 | -5.1 | 19.5 | 47.0 | 281.8 |
| HydrationFreeEnergy_FreeSolv | 19392 | -7.3 | 3.1 | -21.0 | -8.9 | -6.9 | -5.1 | 0.4 |
| LD50_Zhu | 19392 | 3.0 | 0.7 | 1.2 | 2.5 | 3.0 | 3.6 | 5.4 |
| Lipophilicity_AstraZeneca | 19392 | 3.0 | 1.7 | -4.1 | 2.1 | 3.4 | 4.3 | 6.1 |
| PPBR_AZ | 19392 | 95.6 | 18.4 | -24.6 | 87.6 | 99.9 | 107.7 | 123.6 |
| Solubility_AqSolDB | 19392 | -5.1 | 2.1 | -9.2 | -6.7 | -5.5 | -3.6 | 1.5 |
| VDss_Lombardo | 19392 | 4.5 | 5.7 | -14.7 | 0.8 | 3.8 | 7.5 | 47.4 |

**SI Table 5:** Literature evidence for the transformation products.

| Parent | CID (Parent) | Product | CID (Product) | Transformation type | PubMed Link |
| --- | --- | --- | --- | --- | --- |
| Milbemectin | 9828343 | 27-Hydroxy milbemectin A3 | 76334273 | Hydroxylation | [12596855](https://pubmed.ncbi.nlm.nih.gov/12596855) |
| Milbemectin | 9828343 | 27-oxomilbemycin A3 | 76319811 | Oxidation | [12596855](https://pubmed.ncbi.nlm.nih.gov/12596855) |
| (E)-1,3-Dichloropropene | 24726 | 2-Propen-1-ol, 3-chloro-, (2E)- | 287955 | Hydrolysis | [9687453](https://pubmed.ncbi.nlm.nih.gov/9687453) |
| (E)-1,3-Dichloropropene | 24726 | cis-3-Chloroallyl alcohol | 643775 | Hydrolysis | [9687453](https://pubmed.ncbi.nlm.nih.gov/9687453) |
| (E)-1,3-Dichloropropene | 24726 | 2,3-Dichloropropionaldehyde | 93058 | Oxidation | [16841964](https://pubmed.ncbi.nlm.nih.gov/16841964) |
| (E)-Methomyl | 5353758 | Methomyl oxime | 5365286 | Hydrolysis | [28083911](https://pubmed.ncbi.nlm.nih.gov/28083911) |
| Clofenotane | 3036 | 2,3-Dihydroxy-DDT | 441118 | dihydroxylation | [15018096](https://pubmed.ncbi.nlm.nih.gov/15018096) |
| Clofenotane | 3036 | (1S,2S)-DDT-2,3-dihydrodiol | 441117 | Oxidation | [8117093](https://pubmed.ncbi.nlm.nih.gov/8117093) |
| Acephate | 1982 | Methamidophos | 4096 | Hydrolysis | [9074804](https://pubmed.ncbi.nlm.nih.gov/9074804) |
| Acequinocyl | 93315 | 2-Dodecyl-3-hydroxy-1,4-naphthoquinone | 14672776 | Hydrolysis | [15506803](https://pubmed.ncbi.nlm.nih.gov/15506803) |
| Aldicarb | 9570071 | Aldoxycarb | 9570093 | Oxidation | [16751141](https://pubmed.ncbi.nlm.nih.gov/16751141) |
| Allethrins | 11442 | 2-Cyclopenten-1-one, 4-hydroxy-3-methyl-2-(2-propen-1-yl)- | 11083 | Hydrolysis | [16328989](https://pubmed.ncbi.nlm.nih.gov/16328989) |
| Azinphos-Methyl | 2268 | Azinphosmethyl oxon | 13746 | Oxidative desulfuration | [11298499](https://pubmed.ncbi.nlm.nih.gov/11298499) |
| Carbaryl | 6129 | 1-Naphthol | 7005 | Hydrolysis | [12929849](https://pubmed.ncbi.nlm.nih.gov/12929849) |
| Carbaryl | 6129 | Carbamic acid, (hydroxymethyl)-, 1-naphthalenyl ester | 21341 | hydroxylation | [6422454](https://pubmed.ncbi.nlm.nih.gov/6422454) |
| Carbaryl | 6129 | 1-naphthalenyl N-methyl-N-nitrosocarbamate | 23493 | Nitrosation | [6785222](https://pubmed.ncbi.nlm.nih.gov/6785222) |
| Carbofuran | 2566 | 2,2-Dimethyl-2,3-dihydro-1-benzofuran-4,7-diol | 15175289 | Oxidation | [12351226](https://pubmed.ncbi.nlm.nih.gov/12351226) |
| Carbofuran | 2566 | 2,3-Dihydro-2,2-dimethyl-7-benzofuranol | 15278 | Hydrolysis | [17305136](https://pubmed.ncbi.nlm.nih.gov/17305136) |
| Carbofuran | 2566 | 5-Hydroxycarbofuran | 25210295 | Hydroxylation | [17425661](https://pubmed.ncbi.nlm.nih.gov/17425661) |
| Carbofuran | 2566 | 3-Hydroxycarbofuran | 27975 | hydroxylation | [16472538](https://pubmed.ncbi.nlm.nih.gov/16472538) |
| Carbofuran | 2566 | N-Nitrosocarbofuran | 44098 | N-nitrosation | [16089328](https://pubmed.ncbi.nlm.nih.gov/16089328) |
| Carbofuran | 2566 | 3-Ketocarbofuran | 27999 | Oxidation | [17241092](https://pubmed.ncbi.nlm.nih.gov/17241092) |
| Chlordane, technical | 5993 | Chlordene | 19519 | dechlorination | [21718811](https://pubmed.ncbi.nlm.nih.gov/21718811) |
| Chlorpyrifos | 2730 | 3,5,6-Trichloro-2-pyridinol | 23017 | Hydrolysis | [11353146](https://pubmed.ncbi.nlm.nih.gov/11353146) |
| Chlorpyrifos | 2730 | Chlorpyrifos oxon | 21804 | Oxidative desulfuration | [41129308](https://pubmed.ncbi.nlm.nih.gov/41129308) |
| Chlorpyrifos-methyl | 21803 | 3,5,6-Trichloro-2-pyridinol | 23017 | Hydrolysis | [22425279](https://pubmed.ncbi.nlm.nih.gov/22425279) |
| Coumaphos | 2871 | Chlorferon | 5355079 | Hydrolysis | [20134347](https://pubmed.ncbi.nlm.nih.gov/20134347) |
| Coumaphos | 2871 | Coumaphos oxon | 9453 | Oxidation | [18826034](https://pubmed.ncbi.nlm.nih.gov/18826034) |
| Cyfluthrin | 104926 | 2-(4-Fluoro-3-phenoxyphenyl)-2-hydroxyacetonitrile | 10377109 | Hydrolysis | [15865349](https://pubmed.ncbi.nlm.nih.gov/15865349) |
| Cypermethrin | 2912 | m-Phenoxybenzyl cyanide | 40075 | ester cleavage | [11852644](https://pubmed.ncbi.nlm.nih.gov/11852644) |
| Cypermethrin | 2912 | Benzeneacetonitrile, alpha-hydroxy-3-phenoxy-, (alphaS)- | 93218 | Hydrolysis | [22038248](https://pubmed.ncbi.nlm.nih.gov/22038248) |
| Deltamethrin | 40585 | m-Phenoxybenzyl cyanide | 40075 | Hydrolysis | [17576809](https://pubmed.ncbi.nlm.nih.gov/17576809) |
| Diazinon | 3017 | 2-Isopropyl-6-methyl-4-pyrimidinol | 135444498 | Hydrolysis | [18437620](https://pubmed.ncbi.nlm.nih.gov/18437620) |
| Diazinon | 3017 | O,O-Diethyl O-(2-(1-hydroxy-1-methylethyl)-6-methylpyrimidin-4-yl)phosphorothioate | 592775 | Hydroxylation | [21733545](https://pubmed.ncbi.nlm.nih.gov/21733545) |
| Diazinon | 3017 | Diazoxon | 13754 | Oxidation | [17132760](https://pubmed.ncbi.nlm.nih.gov/17132760) |
| Dichlorvos | 3039 | Demethyl dichlorvos | 176943 | demethylation | [17937289](https://pubmed.ncbi.nlm.nih.gov/17937289) |
| Dicofol | 8268 | 4,4'-Dichloro-alpha-(dichloromethyl)benzhydrol | 160701 | Reductive dechlorination | [3424865](https://pubmed.ncbi.nlm.nih.gov/3424865) |
| Dimethoate | 3082 | Dimethyl phosphorothioate | 168140 | Hydrolysis | [22884955](https://pubmed.ncbi.nlm.nih.gov/22884955) |
| Dimethoate | 3082 | O-Demethyldimethoate | 3080588 | O-demethylation | [6658030](https://pubmed.ncbi.nlm.nih.gov/6658030) |
| Dimethoate | 3082 | Omethoate | 14210 | Oxidation | [6736564](https://pubmed.ncbi.nlm.nih.gov/6736564) |
| Disulfoton | 3118 | Oxydisulfoton | 17242 | Oxidation | [22818267](https://pubmed.ncbi.nlm.nih.gov/22818267) |
| Disulfoton | 3118 | Disulfoton sulfone | 17241 | Oxidation | [22818267](https://pubmed.ncbi.nlm.nih.gov/22818267) |
| Endosulfan | 3224 | Endosulfan Sulfate | 13940 | Oxidation | [21550631](https://pubmed.ncbi.nlm.nih.gov/21550631) |
| Etoxazole | 153974 | 2-Amino-2-(4-tert-butyl-2-ethoxyphenyl)ethanol | 15346092 | Hydrolysis | [21667235](https://pubmed.ncbi.nlm.nih.gov/21667235) |
| Fenamiphos | 31070 | Fenamiphos sulfoxide | 36027 | Oxidation | [27554030](https://pubmed.ncbi.nlm.nih.gov/27554030) |
| Fenamiphos | 31070 | Fenamiphos sulfone | 36028 | Oxidation | [27554030](https://pubmed.ncbi.nlm.nih.gov/27554030) |
| Fenitrothion | 31200 | 3-Methyl-4-nitrophenol | 17412 | Hydrolysis | [17625231](https://pubmed.ncbi.nlm.nih.gov/17625231) |
| Fenitrothion | 31200 | Carboxyfenitrothion | 41194 | Oxidation | [17822740](https://pubmed.ncbi.nlm.nih.gov/17822740) |
| Fenitrothion | 31200 | Fenitrooxon | 16738 | Oxidation | [24038429](https://pubmed.ncbi.nlm.nih.gov/24038429) |
| Fenitrothion | 31200 | Aminofenitrothion | 25843 | Reduction | [3824409](https://pubmed.ncbi.nlm.nih.gov/3824409) |
| Fenitrothion | 31200 | 4-Nitrosofenitrothion | 127435 | Reduction | [3824409](https://pubmed.ncbi.nlm.nih.gov/3824409) |
| Fenobucarb | 19588 | 2-sec-Butylphenol | 6984 | Hydrolysis | [24197220](https://pubmed.ncbi.nlm.nih.gov/24197220) |
| Fenpropathrin | 47326 | Benzeneacetonitrile, alpha-hydroxy-3-phenoxy-, (alphaS)- | 93218 | Hydrolysis | [21727000](https://pubmed.ncbi.nlm.nih.gov/21727000) |
| Fenthion | 3346 | 3-Methyl-4-(methylthio)phenol | 18391 | Photolysis | [20022167](https://pubmed.ncbi.nlm.nih.gov/20022167) |
| Fenvalerate | 3347 | Benzeneacetonitrile, alpha-hydroxy-3-phenoxy-, (alphaS)- | 93218 | Hydrolysis | [21727000](https://pubmed.ncbi.nlm.nih.gov/21727000) |
| Fenvalerate | 3347 | m-Phenoxybenzyl cyanide | 40075 | Hydrolysis | [17576809](https://pubmed.ncbi.nlm.nih.gov/17576809) |
| Fipronil | 3352 | Fipronil sulfone | 3078139 | Oxidation | [28343720](https://pubmed.ncbi.nlm.nih.gov/28343720) |
| Flonicamid | 9834513 | N-(4-Trifluoromethylnicotinoyl)glycine | 46835486 | Hydrolysis | [23016287](https://pubmed.ncbi.nlm.nih.gov/23016287) |
| Flonicamid | 9834513 | 3-Pyridinecarboxylic acid, 4-(trifluoromethyl)- | 2777549 | Hydrolysis | [23016287](https://pubmed.ncbi.nlm.nih.gov/23016287) |
| Flonicamid | 9834513 | 4-Trifluoromethylnicotinamide | 2782643 | N-dealkylation | [23016287](https://pubmed.ncbi.nlm.nih.gov/23016287) |
| Hexachlorocyclohexane | 727 | gamma-Pentachlorocyclohexene | 441093 | dehydrochlorination | [7686793](https://pubmed.ncbi.nlm.nih.gov/7686793) |
| Hexachlorocyclohexane | 727 | 3,4,5,6-Tetrachlorocyclohexene | 15691 | dichloroelimination | [16202806](https://pubmed.ncbi.nlm.nih.gov/16202806) |
| Imidacloprid | 86287518 | Desnitro-imidacloprid | 10130527 | Denitration | [34628512](https://pubmed.ncbi.nlm.nih.gov/34628512) |
| Imidacloprid | 86287518 | Imidacloprid urea | 15390532 | Hydrolysis | [18690690](https://pubmed.ncbi.nlm.nih.gov/18690690) |
| Imidacloprid | 86287518 | [C(E)]-N-[(6-Chloro-3-pyridinyl)methyl]-N'-nitro-guanidine | 135833314 | Oxidation | [15849408](https://pubmed.ncbi.nlm.nih.gov/15849408) |
| Isoprocarb | 17517 | 2-Isopropylphenol | 6943 | Hydrolysis | [19071416](https://pubmed.ncbi.nlm.nih.gov/19071416) |
| Malathion | 4004 | Diethyl 2-[hydroxy(methoxy)phosphinothioyl]sulfanylbutanedioate | 50904182 | Hydrolysis | [5950619](https://pubmed.ncbi.nlm.nih.gov/5950619) |
| Malathion | 4004 | Monoethyl ((dimethoxyphosphinothioyl)thio)butanedioate | 20337862 | Hydrolysis | [402103](https://pubmed.ncbi.nlm.nih.gov/402103) |
| Malathion | 4004 | Malaoxon | 15415 | Oxidation | [8292751](https://pubmed.ncbi.nlm.nih.gov/8292751) |
| Methamidophos | 4096 | O,S-Dimethyl hydrogen phosphorothioate | 23413258 | Hydrolysis | [19960233](https://pubmed.ncbi.nlm.nih.gov/19960233) |
| Methiocarb | 16248 | Methiocarb sulfoxide | 17521 | Oxidation | [23252625](https://pubmed.ncbi.nlm.nih.gov/23252625) |
| Methoxychlor | 4115 | Methoxychlor olefin | 75048 | dehydrochlorination | [16753958](https://pubmed.ncbi.nlm.nih.gov/16753958) |
| Methoxychlor | 4115 | Monodemethylmethoxychlor | 183679 | O-demethylation | [20840852](https://pubmed.ncbi.nlm.nih.gov/20840852) |
| Methoxychlor | 4115 | 1,1'-(2,2-Dichloroethylidene)bis[4-methoxybenzene] | 81869 | Reductive dechlorination | [17906828](https://pubmed.ncbi.nlm.nih.gov/17906828) |
| Methyl Parathion | 4130 | 4-Nitrophenol | 980 | Hydrolysis | [11701217](https://pubmed.ncbi.nlm.nih.gov/11701217) |
| Methyl Parathion | 4130 | Paraoxon methyl | 13708 | Oxidation | [19539977](https://pubmed.ncbi.nlm.nih.gov/19539977) |
| Methyl Parathion | 4130 | O-(4-Aminophenyl) O,O-dimethyl phosphorothioate | 44152294 | Reduction | [14758519](https://pubmed.ncbi.nlm.nih.gov/14758519) |
| Methyl Parathion | 4130 | Dimethoxy-(4-nitrosophenoxy)-sulfanylidene-lambda5-phosphane | 85992250 | Reduction | [14758519](https://pubmed.ncbi.nlm.nih.gov/14758519) |
| Metolcarb | 14322 | M-Cresol | 342 | Hydrolysis | [19071416](https://pubmed.ncbi.nlm.nih.gov/19071416) |
| Metolcarb | 14322 | N-ME-3-Formylphenylcarbamate | 235375 | Oxidation | [894678](https://pubmed.ncbi.nlm.nih.gov/894678) |
| Acetamiprid | 213021 | (E)-N-((6-chloropyridin-3-yl)methyl)-N'-cyanoethanimidamide | 11344811 | N-demethylation | [21713509](https://pubmed.ncbi.nlm.nih.gov/21713509) |
| Naled | 4420 | Dichlorvos | 3039 | debromination | [36912955](https://pubmed.ncbi.nlm.nih.gov/36912955) |
| Nicotine | 89594 | nicotine N-glucuronide | 71751013 | Glucuronidation | [36252563](https://pubmed.ncbi.nlm.nih.gov/36252563) |
| Nicotine | 89594 | (S)-6-Hydroxynicotine | 439383 | hydroxylation | [4405081](https://pubmed.ncbi.nlm.nih.gov/4405081) |
| Nicotine | 89594 | (3S,5S)-1-methyl-5-pyridin-3-ylpyrrolidin-3-ol | 12960088 | Hydroxylation | [7932582](https://pubmed.ncbi.nlm.nih.gov/7932582) |
| Nicotine | 89594 | Nornicotine | 91462 | N-demethylation | [4053005](https://pubmed.ncbi.nlm.nih.gov/4053005) |
| Nicotine | 89594 | 4-(Methylnitrosamino)-4-(3-pyridyl)butyric acid | 115108 | Nitrosation | [8330358](https://pubmed.ncbi.nlm.nih.gov/8330358) |
| Nicotine | 89594 | 4-(Methylnitrosamino)-4-(3-pyridyl)butanal | 62228 | Nitrosation | [633391](https://pubmed.ncbi.nlm.nih.gov/633391) |
| Nicotine | 89594 | Nicotine 1'-N-oxide | 68107 | N-oxidation | [16536909](https://pubmed.ncbi.nlm.nih.gov/16536909) |
| Nicotine | 89594 | Cotinine | 854019 | Oxidation | [21208832](https://pubmed.ncbi.nlm.nih.gov/21208832) |
| Nicotine | 89594 | (2S)-2-(Pyridin-3-yl)pyrrolidine-1-carbaldehyde | 13945370 | Oxidation | [24581918](https://pubmed.ncbi.nlm.nih.gov/24581918) |
| Nicotine | 89594 | nicotine N-oxide | 211457 | Oxidation | [37787014](https://pubmed.ncbi.nlm.nih.gov/37787014) |
| Nicotine | 89594 | 3-Vinylpyridine | 14272 | Oxidation | [29277437](https://pubmed.ncbi.nlm.nih.gov/29277437) |
| Nicotine | 89594 | 4-Hydroxy-4-(3-pyridyl)butanoic acid | 438 | Oxidation | [21452992](https://pubmed.ncbi.nlm.nih.gov/21452992) |
| Nicotine | 89594 | N-Methyl-gamma-oxo-3-pyridinebutanamide | 436 | Oxidation | [7098975](https://pubmed.ncbi.nlm.nih.gov/7098975) |
| Nicotine | 89594 | 4-Oxo-4-(3-pyridyl)butyric acid | 437 | Oxidation | [27567546](https://pubmed.ncbi.nlm.nih.gov/27567546) |
| Nicotine | 89594 | 3-(1-Methyl-1-oxido-2-pyrrolidinyl)pyridine | 409 | Oxidation | [11377240](https://pubmed.ncbi.nlm.nih.gov/11377240) |
| Nicotine | 89594 | Nicotine iminium | 181139 | Oxidation | [30906561](https://pubmed.ncbi.nlm.nih.gov/30906561) |
| Nicotine | 89594 | 3-Hydroxycotinine | 10219774 | Oxidation | [8468378](https://pubmed.ncbi.nlm.nih.gov/8468378) |
| Oxydemeton-methyl | 4618 | Demeton-S-methyl | 13526 | Reduction | [19436153](https://pubmed.ncbi.nlm.nih.gov/19436153) |
| Parathion | 991 | O-Ethyl O-(4-nitrophenyl) phosphorothioate | 21447693 | Hydrolysis | [16209589](https://pubmed.ncbi.nlm.nih.gov/16209589) |
| Parathion | 991 | 4-Nitrophenol | 980 | Hydrolysis | [12669189](https://pubmed.ncbi.nlm.nih.gov/12669189) |
| Parathion | 991 | Paraoxon | 9395 | Oxidation | [27256520](https://pubmed.ncbi.nlm.nih.gov/27256520) |
| Parathion | 991 | Aminoparathion | 220 | Reduction | [16277414](https://pubmed.ncbi.nlm.nih.gov/16277414) |
| Pentachlorophenol | 992 | 2,4,6-Trichlorophenol | 6914 | dechlorination | [8867638](https://pubmed.ncbi.nlm.nih.gov/8867638) |
| Pentachlorophenol | 992 | 3,4,5-Trichlorophenol | 11859 | dechlorination | [29127932](https://pubmed.ncbi.nlm.nih.gov/29127932) |
| Pentachlorophenol | 992 | 3-Chlorophenol | 7933 | dechlorination | [23479749](https://pubmed.ncbi.nlm.nih.gov/23479749) |
| Pentachlorophenol | 992 | Pentachlorophenol glucuronide | 131876757 | Glucuronidation | [3977960](https://pubmed.ncbi.nlm.nih.gov/3977960) |
| Pentachlorophenol | 992 | Tetrachlorocatechol | 14537 | Oxidation | [20452725](https://pubmed.ncbi.nlm.nih.gov/20452725) |
| Pentachlorophenol | 992 | Tetrachloro-p-hydroquinone | 66603 | oxidative dechlorination | [9751251](https://pubmed.ncbi.nlm.nih.gov/9751251) |
| Pentachlorophenol | 992 | Pentachlorobenzene | 11855 | Reduction | [17962225](https://pubmed.ncbi.nlm.nih.gov/17962225) |
| Pentachlorophenol | 992 | 2,3,4,5-Tetrachlorophenol | 21013 | Reductive dechlorination | [12948058](https://pubmed.ncbi.nlm.nih.gov/12948058) |
| Pentachlorophenol | 992 | 2,3,5-Trichlorophenol | 13619 | Reductive dechlorination | [11731034](https://pubmed.ncbi.nlm.nih.gov/11731034) |
| Pentachlorophenol | 992 | 2,3,6-Trichlorophenol | 13618 | Reductive dechlorination | [17438781](https://pubmed.ncbi.nlm.nih.gov/17438781) |
| Pentachlorophenol | 992 | 2,3-Dichlorophenol | 11334 | Reductive dechlorination | [17438781](https://pubmed.ncbi.nlm.nih.gov/17438781) |
| Pentachlorophenol | 992 | 3,5-Dichlorophenol | 11571 | Reductive dechlorination | [22743203](https://pubmed.ncbi.nlm.nih.gov/22743203) |
| Permethrin | 40326 | 3-Phenoxybenzyl alcohol | 26295 | Hydrolysis | [21456540](https://pubmed.ncbi.nlm.nih.gov/21456540) |
| Phenothrin | 4767 | 3-Phenoxybenzyl alcohol | 26295 | Hydrolysis | [31538121](https://pubmed.ncbi.nlm.nih.gov/31538121) |
| Phenthoate | 17435 | Ethyl alpha-((dimethoxyphosphinyl)thio)benzeneacetate | 19398 | Oxidative desulfuration | [6826909](https://pubmed.ncbi.nlm.nih.gov/6826909) |
| Phorate | 4790 | O,O-Diethyl dithiophosphate | 9274 | Hydrolysis | [19413758](https://pubmed.ncbi.nlm.nih.gov/19413758) |
| Phorate | 4790 | Phorate sulfone | 17425 | Oxidation | [19413758](https://pubmed.ncbi.nlm.nih.gov/19413758) |
| Phorate | 4790 | Phorate sulfoxide | 17424 | sulfoxidation | [19413758](https://pubmed.ncbi.nlm.nih.gov/19413758) |
| Pirimicarb | 31645 | Pirimicarb III | 93139 | Hydrolysis | [19268958](https://pubmed.ncbi.nlm.nih.gov/19268958) |
| Pirimicarb | 31645 | Pirimicarb-desamido | 135420611 | Hydrolysis | [20117325](https://pubmed.ncbi.nlm.nih.gov/20117325) |
| Pirimiphos-methyl | 34526 | 2-(diethylamino)-6-methyl-1H-pyrimidin-4-one | 135448147 | Hydrolysis | [17687456](https://pubmed.ncbi.nlm.nih.gov/17687456) |
| Prallethrin | 9839306 | Allethrins | 11442 | Reduction | [30085203](https://pubmed.ncbi.nlm.nih.gov/30085203) |
| Propoxur | 4944 | 2-Isopropoxyphenol | 20949 | Hydrolysis | [17107853](https://pubmed.ncbi.nlm.nih.gov/17107853) |
| Propoxur | 4944 | 2-(1-Methylethoxy)-5-nitrophenyl N-methylcarbamate | 91431723 | Nitration | [21581688](https://pubmed.ncbi.nlm.nih.gov/21581688) |
| Propoxur | 4944 | 2-(1-Methylethoxy)phenyl N-methyl-N-nitrosocarbamate | 38097 | Nitrosation | [18329776](https://pubmed.ncbi.nlm.nih.gov/18329776) |
| Rotenone | 6758 | 12A-Hydroxyrotenone | 68184 | Hydroxylation | [17658825](https://pubmed.ncbi.nlm.nih.gov/17658825) |
| Terbufos | 25670 | Terbufos sulfoxide | 25355 | Oxidation | [20803314](https://pubmed.ncbi.nlm.nih.gov/20803314) |
| Terbufos | 25670 | Terbufos sulfone | 41718 | Oxidation | [22043870](https://pubmed.ncbi.nlm.nih.gov/22043870) |
| Tetramethrin | 83975 | Chrysanthemic Acid | 2743 | Hydrolysis | [16257148](https://pubmed.ncbi.nlm.nih.gov/16257148) |
| Tolfenpyrad | 10110536 | Tolfenpyrad-benzoic acid | 123319859 | Oxidation | [22802573](https://pubmed.ncbi.nlm.nih.gov/22802573) |
| Transfluthrin | 656612 | 2,3,5,6-Tetrafluorobenzyl alcohol | 2734029 | Hydrolysis | [24560337](https://pubmed.ncbi.nlm.nih.gov/24560337) |
| Triazamate | 86306 | Triazamate acid | 14299898 | Hydrolysis | [15204546](https://pubmed.ncbi.nlm.nih.gov/15204546) |
| Triazophos | 32184 | 1-Phenyl-3-hydroxy-1,2,4-triazole | 77910 | Hydrolysis | [21592646](https://pubmed.ncbi.nlm.nih.gov/21592646) |
| Trichlorfon | 5853 | Dimethyl P-(1-(acetyloxy)-2,2,2-trichloroethyl)phosphonate | 22235 | Acetylation | [3857760](https://pubmed.ncbi.nlm.nih.gov/3857760) |
| Trichlorfon | 5853 | Demethyldichlorvos | 114636 | degradation | [7074710](https://pubmed.ncbi.nlm.nih.gov/7074710) |
| Trichlorfon | 5853 | Dichlorvos | 3039 | dehydrochlorination | [1475794](https://pubmed.ncbi.nlm.nih.gov/1475794) |
| Trichlorfon | 5853 | Butonate | 31343 | Esterification | [7074710](https://pubmed.ncbi.nlm.nih.gov/7074710) |
| Trichlorfon | 5853 | Dimethyl phosphate | 13134 | Hydrolysis | [4000249](https://pubmed.ncbi.nlm.nih.gov/4000249) |
| Sulfluramid | 77797 | Perfluorooctanesulfonic acid | 74483 | environmental biotransformation | [29793104](https://pubmed.ncbi.nlm.nih.gov/29793104) |
| Sulfluramid | 77797 | Perfluorooctanesulfonamide | 69785 | N-deethylation | [36914008](https://pubmed.ncbi.nlm.nih.gov/36914008) |

**Supplementary method:**

We codified a robust and systematic workflow for the data-driven discovery and application of chemical transformation rules, designed for predicting metabolic fate, environmental degradation, or synthetic outcomes. This github repo <https://github.com/idslme/in-silico-transformation> contains all the scripts to describe the workflow. The workflow is created as a sequential pipeline of seven Python scripts, each performing a dedicated task, transforming raw data from the PubChem repository into a predictive engine.

1_pubchem_rxns_compilation.py : This initial script serves as the data acquisition module, systematically harvesting raw reaction data from the PubChem database via its Power User Gateway (PUG) API. Its operation is not a blind download but a targeted query.

2_rxn_equ_format_conversion.py: This script is the standardization and translation layer. Raw CIDs are abstract identifiers; this script translates them into a chemically-intelligible, structure-based format required by all downstream cheminformatics tools. The script reads the CID-based dataset from Script 1. For each reactant and product CID, it must again query the PUG API, this time to fetch the canonical SMILES (Simplified Molecular-Input Line-Entry System) string for each compound. It then assembles these individual SMILES into a single, complete reaction SMILES string (e.g., reactant1.reactant2>>product1.product2).

3_rxn_info_cli.py & 4_rxn_info_batch_processing.py : These scripts perform high-throughput validation, feature extraction, and preprocessing on the standardized reaction SMILES. Script 3 is likely a command-line utility for single-reaction inspection, while Script 4 is the parallelized, production-scale version designed for processing massive datasets. Script 4 module is critical for ensuring data quality and preparing reactions for the complex rule-extraction step. It initializes RDKit reaction objects from the SMILES strings and performs several key operations. It checks for chemical validity and stoichiometric balance. It attempts to establish atom-atom mapping (AAM), which is *essential* for template extraction. This process assigns a unique index to each atom, tracking its position from reactant to product. It differentiates between reactants, products, and agents (catalysts, solvents) using RDKit's reaction object properties.

5_reaction_template_generation.py: This is the main logical core of the entire workflow: the inductive rule-generation module. It takes the atom-mapped reactions and generalizes them into abstract transformation templates, or "rules." This script leverages the rdchiral library, a tool specifically designed for extracting reaction templates from atom-mapped reactions. For each reaction, rdchiral identifies the reaction center—the collection of atoms and bonds that are broken, formed, or changed (e.g., in charge or aromaticity). It then generates a generalized SMARTS (SMiles ARbitrary Target Specification) pattern that represents this transformation, ignoring the unchanged parts of the molecules.

6_substructure_generation.py : This script performs site-of-reactivity (SoR) analysis, adding crucial chemical context to the abstract templates. A template (e.g., "aromatic hydroxylation") is a rule, but this script investigates *why* that rule applied to a *specific location* (a specific carbon atom) on the reactant. This module identifies the "metabolic hotspots." It takes a reactant molecule and the template that transformed it. Using the template's reactant-side SMARTS, it performs a substructure match on the reactant molecule to identify the *exact* atoms involved (the reaction center). It then generates a substructure fragment representing the local chemical environment around that center (e.g., all atoms within a 2- or 3-bond radius).

7_biotransformation.py : This final script is the predictive engine of the entire workflow. It operationalizes the generated knowledge to perform *in silico* screening. This script takes the database of transformation rules from Script 5 (--templates <transformation_templates.csv>) and applies them in a "forward" direction to a list of novel molecules from --input <input_compounds.csv>. For each input compound, it iterates through every template in the database. It again uses rdchiral to perform this forward synthesis.
